# Photoactivatable Plasma Membrane Probe Through Self-Triggered Photooxidation Cascade for Live Super-Resolution Microscopy

**DOI:** 10.1101/2024.05.28.596159

**Authors:** Sonia Pfister, Valentine Le Berruyer, Kyong Fam, Mayeul Collot

## Abstract

Super-resolution imaging based on the localization of single emitters requires a spatio-temporal control of the ON and OFF state. To this end, photoactivatable fluorophores are adapted as they can be turned on upon light irradiation. Here we present a concept called Self-Triggered Photooxidation Cascade (STPC) based on the photooxidation of a plasma membrane targeted leuco-rhodamine (LRhod-PM), a non-fluorescent reduced form of a rhodamine probe. Upon visible light irradiation the small number of oxidized rhodamines, Rhod-PM, acts as a photosensitizer to generate singlet oxygen capable to oxidize the OFF state LRhod-PM. We showed that this phenomenon is kinetically favored by a high local concentration and propagates quickly when the probe is embedded in membrane bilayers. In addition, we showed that the close proximity of the dyes favors the photobleaching. At the single-molecule level, the concomitant activation/bleaching phenomena allow reaching a single-molecule blinking regime enabling single-molecule localization microscopy for super-resolution of live cellular membranes.

## Introduction

Photomodulation of fluorescent molecules draw a particular attention due to their application in advanced bioimaging. Among them, photoactivatable probes are of particular interest in single-molecule localization microscopy to break the diffraction limit and reach super-resolution imaging.^[1], [2], [3], [4], [5], [6], [7]^ Although photoactivatable fluorescent proteins have been developed for this endeavor,^[8], [9]^ the latter remain limited to protein labeling and by their brightness. Consequently, small fluorescent photoactivatable probes constitute an appealing alternative to proteins. In the last decades, several mechanisms have been proposed to develop photoactivatable fluorescent probes. Among other mechanisms,^[10]^ two main different photoactivation processes can be distinguished. 1) Those that involve a photo elimination step including: a nitrogen (N_2_) from a diazoketone^[11]^ or azide-containing fluorophores,^[3], [12]^ elimination of nitric oxide (NO),^[13]^ photocleavage of spirooxazime,^[6]^ and photo-uncaging of fluorescence quenchers.^[14], [15], [16], [17]^ And 2) those involving an addition step, including protonation,^[18], [19]^ and addition of oxygen through a photooxidation process.^[20], [21], [22]^ In this context, we recently established a new mechanism allowing to obtain photoconvertible probes based on directed photooxidation,^[23], [24]^ which enabled live single-molecule localization microscopy (SMLM) of mitochondria and plasma membrane (PM) on fragile samples like neurons.^[25]^ Despite this last example, photoactivatable probes have rarely been targeted to the plasma membrane to perform live SMLM based on PhotoActivated Localization Microscopy (PALM). Indeed, there is only limited examples of SMLM performed at the plasma membrane using small fluorescent probes.^[26]^ On fixed samples, several groups successfully resolved the PM.^[27], [28]^ We notably reported that PM could be imaged at nanoscopic scale on fixed neurons in two-color 3D-STORM using MemBright probes (MB-Cy3.5).^[29]^ However, live SMLM on cell surface is a challenging task due to the dynamic nature of the plasma membrane. Point Accumulation for Imaging in Nanoscale Topography (PAINT) approaches have successfully been applied to live SMLM of the PM, based on the transient binding of fluorogenic probes including solvatochromic Nile Red and self-quenched dimers with tuned lipophilicity.^[30], [31]^ Additionally, BODIPYs showed interesting blinking properties based on PAINT,^[32]^ or transient formation of red-shifted *J*-aggregates.^[33]^ Finally, live SMLM of *E. coli* membrane has been performed using original rhodamine and silicon rhodamine.^[34]^

As referred above, rhodamine is an interesting scaffold to develop photoactivatable probes for live SMLM.^[18], [20]^ Another potential way to trigger rhodamine fluorescence for PALM microscopy is to recover it from a non-emissive reduced leuco-rhodamine, also called dihydroxyrhodamine, that can be oxidized into its fluorescent rhodamine form. Indeed, dihydroxyxanthene scaffolds like DCHF (2’,7’-dichlorodihydrofluorescein) and DHR123 (dihydroxyrhodamine 123) are widely used to detect ROS.^[35]^ Interestingly, there are some concerns about the use of these ROS detectors, as they have been shown to provoke false positive while detection of ROS, due to their tendency to turn on upon “auto-oxidation” or photooxidation.^[36], [37]^ However, by taking advantage of this photooxidation drawback, leuco-rhodamine dyes emerge as a promising scaffold to be used in PALM microscopy. The involvement of leuco-forms of cationic dyes like ATTO,^[38]^ methylene blue,^[39]^ has already been shown in dSTORM experiments on fixed samples.^[40]^ Although the use of reduced fluorophore has already been shown to enhance single-molecule localization density for live super-resolution imaging, the use of exogeneous, non-physiological sodium borohydride was required for super-resolution imaging.^[41]^ Interestingly, Zhang *et al*. showed that dihydro silicon rhodamine could oxidize upon irradiation and used this feature for live SMLM in mitochondria.^[42]^ In this work, we rationalized the use of leuco-rhodamine in live SMLM. We herein hypothesized that when targeted at the plasma membrane and upon irradiation, the small number of already oxidized rhodamines act as mild photosensitizers, able to oxidize their neighboring leuco-rhodamines in a concentration-dependent manner. This mechanism called Self-Triggered Photooxidation Cascade results in a succession of photoactivation and photobleaching which is a desirable feature for live SMLM.

## Results and Discussion

### Design & Synthesis

The design of our photoactivatable probe is based on several concepts. The first one relies on the fact that the leuco-Rhodamine (LRhod), the reduced form of a rhodamine is no longer fluorescent (OFF state) due to the disruption of the conjugation between the two phenyl rings of the xanthene moiety (Figure 1A). However, in oxidizing conditions this LRhod can be reversibly oxidized into fluorescent rhodamine (ON state), Rhod. The second concept relies on the fact that, upon excitation, a fluorophore can generate singlet oxygen (^1^O_2_) through its excited triplet state. Consequently, we hypothesized that LRhod could get oxidized by a neighboring rhodamine which acts as a photosensitizer (Figure 1A). To favor this mechanism, which relies on diffusion, we targeted LRhod to the plasma membrane. At the initial state, a few numbers of Rhod-PM are already present at the cell surface, due to surrounding ambient oxygen (Figure 1B). However, in the plasma membrane (PM) the probes freely diffuse, which favors the interaction between the photoactivatable probe LRhod-PM and the singlet oxygen generated by its oxidized cognate photosensitizer Rhod-PM. This interaction results in the photoactivation of LRhod-PM and since the ^1^O_2_ generated can be responsible for the photobleaching,^[43]^ it can simultaneously lead to photobleaching of the emissive form (Figure 1C).

**Figure 1.**
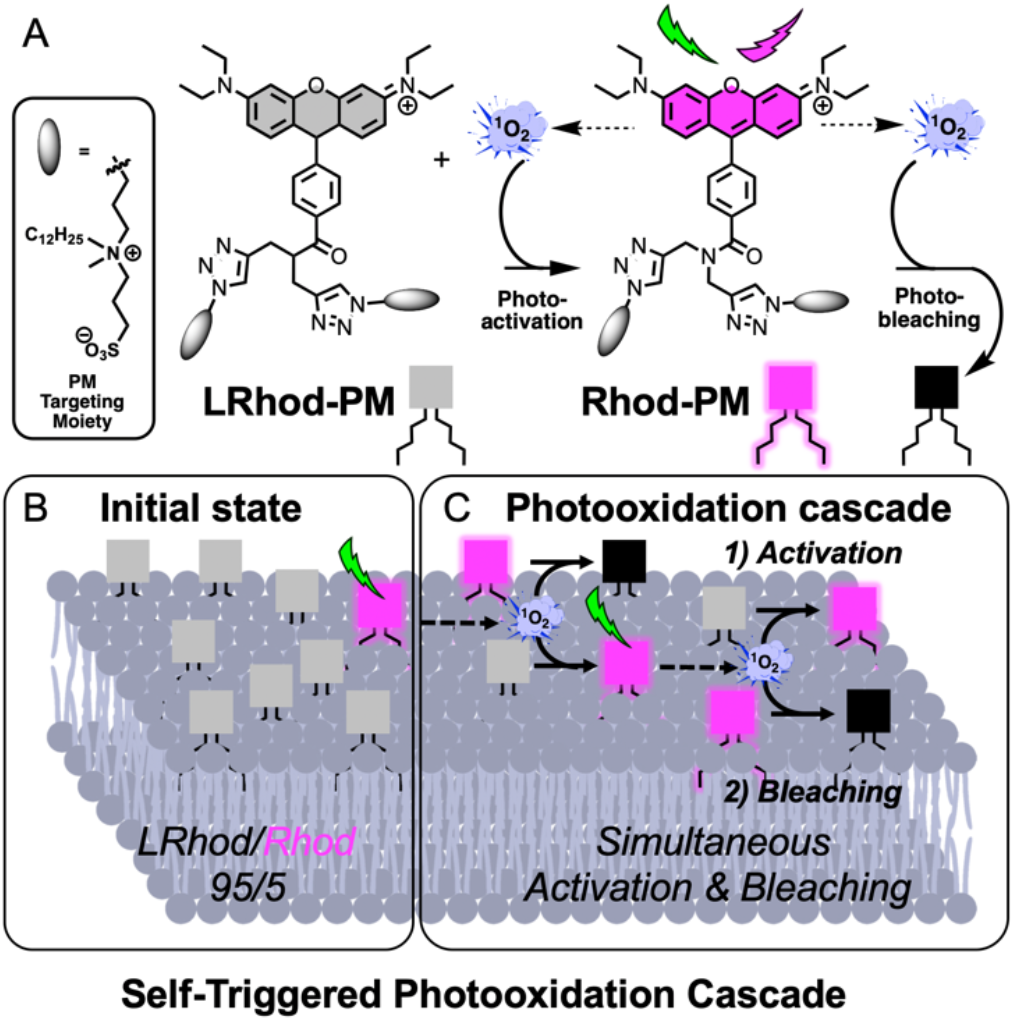
Principle of the Self-Triggered Photooxidation Cascade mechanism. (A) Structures of photoactivatable plasma membrane probe LRhod-PM and its oxidized form Rhod-PM. The principle is based on the ability of Rhod-PM, once concentrated at the membrane, to generate ^1^O_2_ that can oxidize LRhod-PM to generate the fluorescent probe Rhod-PM as well as photobleaching it. (B) At the initial state, the membrane contains a majority of non-emissive LRhod-PM and a small number of Rhod-PM (C) Upon excitation, Rhod-PM acts as a photosensitizer to generate ^1^O_2_ that 1) activates Rhod-PM through oxidation of the neighboring LRhod-PM and 2) provokes its photobleaching.

Overall, we thus expected that an efficient blinking system could be obtained in live SMLM imaging as LRhod-PM probe alternatively turns ON and OFF respectively by photoactivation and photobleaching through a Self-Triggered Photooxidation Cascade (STPC) mechanism.

**Scheme 1.**
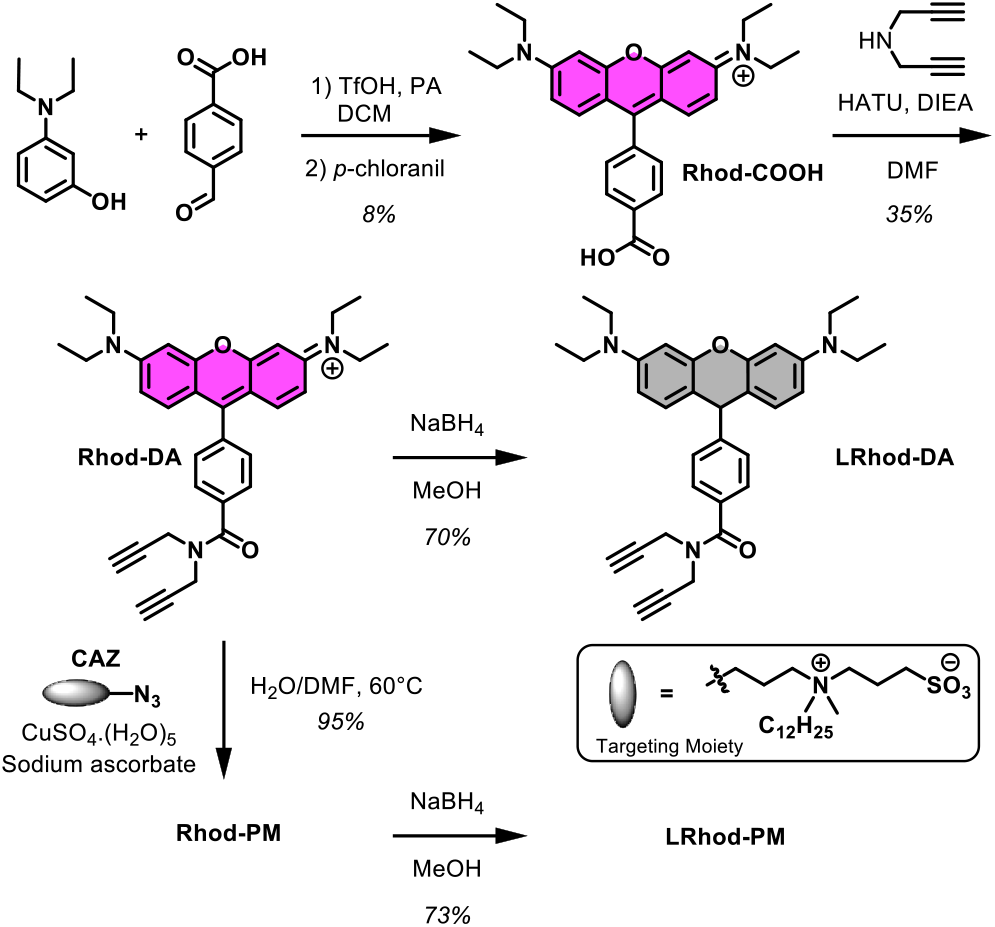
Synthesis of the plasma membrane probes Rhod-PM and LRhod-PM. PA stands for propionic acid, CAZ for Clickable Amphiphilic Zwitterion and DA for DiAlkyne.

To obtain the PM probes, we first synthesized a functionnalizable tetramethyl rhodamine Rhod-COOH possessing a carboxylate group (Scheme 1). The latter was conjugated to dipropargylamine to obtain Rhod-DA which was either reduced by sodium borohydride to provide LRhod-DA or clicked to Clickable Amphiphilic Zwitterion (CAZ) to get Rhod-PM, an approach giving rise to efficient fluorescent plasma membrane probes.^[44], [29], [45], [46], [47], [48]^ As a final step, Rhod-PM was reduced with sodium borohydrate into the non-fluorescent plasma membrane probe leuco-rhodamine LRhod-PM.

### Photophysical Properties

First, the photophysical properties of Rhod-PM and its precursors Rhod-DA and LRhod-DA were evaluated by spectroscopy and are reported in figure 2. As expected, LRhod-DA did not absorb nor emit in the visible range (Figure 2A and B), while Rhod-DA displayed typical maximal excitation and emission wavelengths in methanol (560 and 585 nm respectively) with a bright emission due to a high extinction coefficient (88,200 M^-1^.cm^-1^) and a reasonable quantum yield of fluorescence (ϕ_F_ = 0.23). Rhod-PM rapidly binds to 1,2-dioleoyl-sn-glycero-3-phosphocholine (DOPC) Large Unilamellar Vesicles (LUVs) (Figure S1) and, once embedded into this PM model, displayed slightly red-shifted spectra (Figure 2A) along with slightly reduced brightness (Figure 2B).

**Figure 2.**
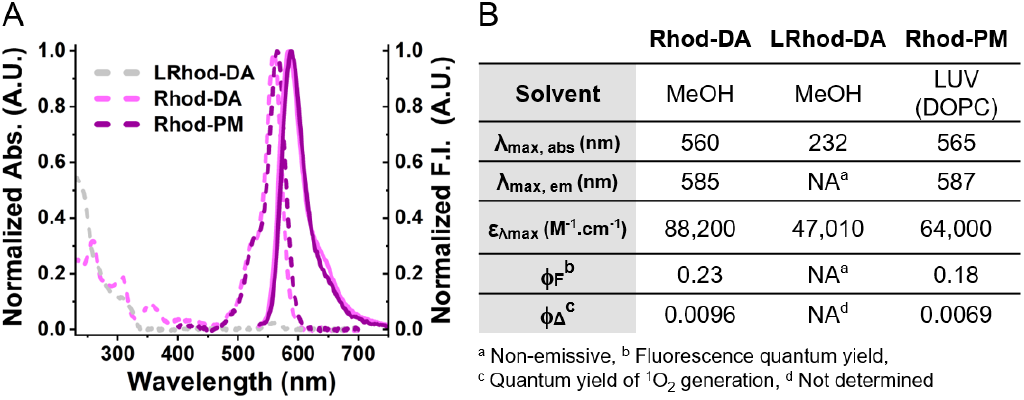
Photophysical properties of the fluorophores. (A) Normalized absorption (dashed lines) and emission (solid lines) spectra of Rhod-PM, Rhod-DA and its reduced form LRhod-DA in MeOH (1 µM). Rhod-PM (1 µM) was embedded into DOPC large unilamellar vesicles (lipid concentration: 200 µM). (B) Photophysical properties of Rhod-DA, LRhod-DA and Rhod-PM.

### Self-Triggered Photooxidation Cascade in Bulk

To prove that a leuco-rhodamine can be photoactivated by a small amount of rhodamine following a self-triggered photooxidation cascade, we first assessed the ability of Rhod-DA to generate singlet oxygen as an oxidizing agent capable of oxidizing a LRhod-DA. Since we expected the rhodamine to act as a photosensitizer, the quantum yields of singlet oxygen generation (ϕ_Δ_) of Rhod-DA and Rhod-PM were evaluated through comparison with rose Bengal as a reference^[49]^ (Figure 3A). The result of 0.96 % and 0.69 % (Figure 2B) was in line with the low intersystem crossing quantum yields of rhodamines (0.5 to 1.0 %),^[50]^ and indicated that Rhod-PM could be used as a mild photosensitizer potentially able to oxidize LRhod-PM without being phototoxic to cells. Then the ability of ^1^O_2_ to oxidize LRhod-DA was assessed by spectroscopy (Figure 3B). LRhod-DA was placed in a methanolic solution in the presence of ^1^O_2_ and the appearance of Rhod-DA was monitored by fluorescence spectroscopy. ^1^O_2_ was generated through the use of methylene blue, an efficient photosensitizer which was independently excited (λ_Ex_= 638 nm, ϕ_Δ_ = 0.50)^[49]^ and which provoked an increase in fluorescence intensity at 585 nm depicting the oxidation of LRhod-DA into its fluorescent oxidized form Rhod-DA. These experiments independently showed that Rhod-DA can be used as a photosensitizer to generate ^1^O_2_ and that the latter is able to oxidize LRhod-DA into its fluorescent Rhod-DA. Therefore, we assumed that Rhod-DA, by itself, could oxidize LRhod-DA. To check our assumption, a concentrated methanolic solution of LRhod-PM (10 µM) was irradiated at 532 nm by a laser beam over time. Upon irradiation, the fluorescence intensity integration increased for 2 hours before slowly decreasing. The plotted curve corresponds to a typical autocatalysis sigmoid profile, relating to a first predominant photoactivation phase followed by a decrease assigned to a prevailing photobleaching phase (Figure 3C). This curve thus suggests 1) that Rhod-DA is generated over time and catalyzed the following oxidation of LRhod-DA and, 2) that upon its formation, Rhod-DA got photobleached. To verify our hypothesis, HPLC studies were conducted in parallel to monitor the photooxidation of LRhod-DA into Rhod-DA (Figure 3D). Before irradiation both LRhod-DA (purple trace) and Rhod-DA (black trace) were analyzed by HPLC and revealed that LRhod-DA contained a small amount of the fluorescent Rhod-DA (≈5%, hashtag # in figure 3D, purple curve), probably due to oxidation over time by the oxygen contained in the air and solvents. Upon irradiation the amount of LRhod-DA decreased (RT= 15.2 min) while Rhod-DA appeared (RT= 22.1 min) along with other photoproducts (*e*.*g*. RT= 21.2 min).

**Figure 3.**
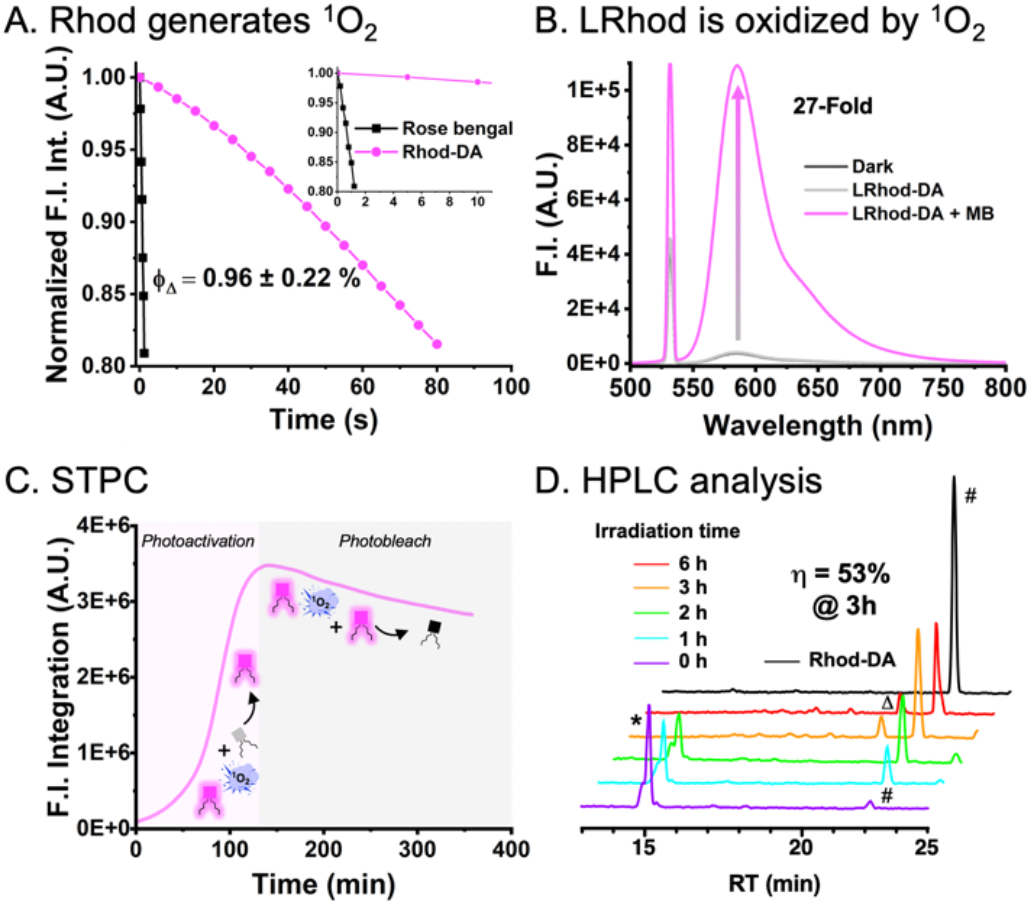
Assessment of the Self-Triggered Photooxidation Cascade (STPC). (A) Singlet oxygen quantum yield measurement: decay of DPBF fluorescence intensity integration in presence of photosensitizers (Rhod-DA or rose bengal as reference) upon 532 nm irradiation. (B) Activation of LRhod-DA (1 µM in methanol) in presence of singlet oxygen sensitizer methylene blue upon 30 minutes of 638 nm irradiation (pink). As control, the same experiment was performed in absence of methylene blue (gray) and in the dark (black curve). (C) Autocatalyzed photo-oxidation of LRhod-DA (10 µM in methanol) over 6 h upon 532 nm irradiation. (D) HPLC traces upon irradiation. The signal was monitored at 254 nm. LRhod-DA and Rhod-DA are respectively indicated by an asterisk and a hashtag. A blue-shifted photoproduct (λ_Abs max_ = 548 nm) is indicated by a delta.

We also observed photoproducts that did not absorb in the visible range depicting a photobleaching whereas others were blue-shifted compared to Rhod-DA (peak indicated by Δ in figure 3D, λ_Abs max_= 548 nm vs 560 nm for Rhod-DA), corresponding to dealkylation upon irradiation.^[51]^ After calibration (Figure S2), we were able to determine a maximum chemical yield of oxidation from LRhod-DA to Rhod-DA of 53% after 3 h of irradiation.

Overall, these analyses confirmed the self-triggered photooxidation cascade process where upon irradiation a small amount of rhodamine acts as a photosensitizer capable tof oxidizing leuco-rhodamine through autocatalysis followed by its photobleaching.

### Self-Triggered Photooxidation Cascade of LRhod-PM in PM Model

After demonstrating that LRhod could be used as a photoactivatable probe, we assessed the self-triggered photooxidation cascade of LRhod-PM at different local concentrations. When LRhod-PM was irradiated in methanol at 200 nM, *i*.*e*. at a low local concentration, virtually no fluorescence enhancement was observed, probably due to the fact that the small number of Rhod-PM present in the bulk was too diluted to trigger the photooxidation cascade of LRhod-PM (Figure 4A). Conversely, when LRhod-PM was incorporated in PM model: DOPC Large Unilamellar Vesicles (LUVs) at a probe/lipid ratio of 1/250, a fluorescence enhancement at 589 nm of 38-fold was observed in 20 min, and a quantum yield of photoactivation was determined, ϕ_Pa_ = 5.8 × 10^−5^ (Figure 4B), which is in the range of photoactivatable fluorescent proteins.^[52], [53]^ This phenomenon was assigned to the increase in local concentration of probes, thus favoring the oxidation of LRhod-PM by its parent photosensitizer Rhod-PM.

**Figure 4.**
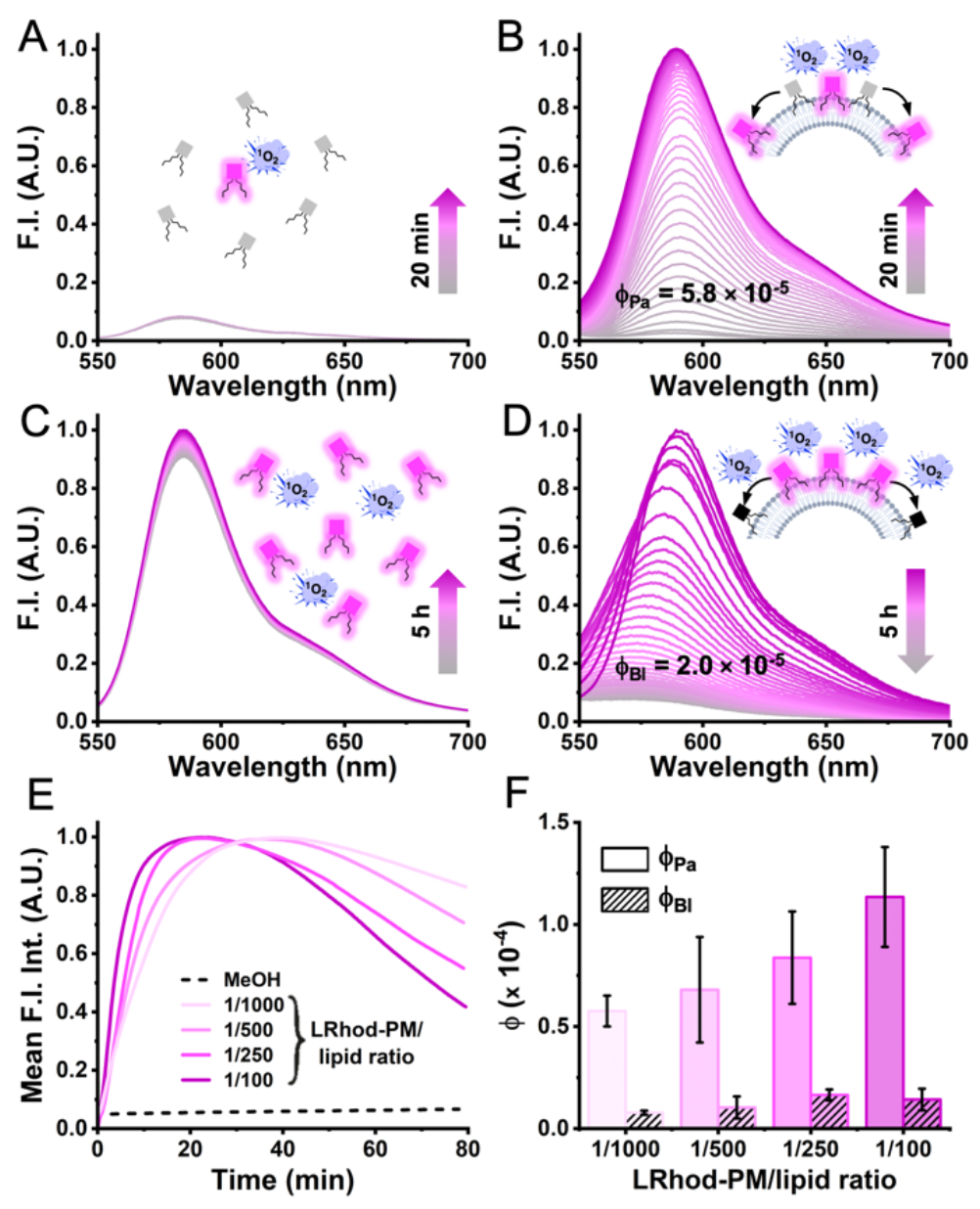
Self-triggered photooxidation cascade of LRhod-PM in low local concentration (200 nM in MeOH) *vs* in high local concentration (200 nM in PBS in presence of LUVs) upon 532 nm irradiation. (A and B) Photoactivation study. Irradiation time: 20 min. (A) In low local concentration and (B) In high local concentration (1/250 probe/lipid). (C and D) Photobleaching study. Irradiation time: 5 h (C) In low local concentration and (D) In high local concentration: (1/250 probe/lipid). (E) Mean fluorescence intensity integration of LRhod-PM in MeOH and in LUVs at different probe/lipid ratios over time upon 532 nm irradiation (200 nM LRhod-PM, probe/lipid ratio: 1/100, 1/250, 1/500, 1/1000). (F) Determination of ϕ_Pa_ and ϕ_Bl_ in different dye/lipids ratios. Three independent experiences were analyzed.

The same trend was observed for the photobleaching phenomenon when the fluorescent Rhod-PM was irradiated in methanol and in LUVs (Figure 4C & D). After 5 h of irradiation in methanol, a slight fluorescence increase was observed assigned to solvent evaporation while due to redistribution of the probe leading to a higher local concentration in LUVs, the kinetic of photobleaching was considerably increased compared to conditions in methanol when the probe homogeneously distributed, with a quantum yield of photobleaching, ϕ_Bl_ of 2.0 × 10^−5^ (Figure 4D). It was noteworthy that upon photobleaching in the LUVs a significant blue shift was observed which is in line with our previous HPLC analysis and the photo-dealkylation process.^[51]^ Consequently, when localized in a lipid membrane, LRhod-PM alternatively undergoes enhanced photoactivation and enhanced photobleaching through a self-triggered photooxidation cascade due to increased local concentration. This hypothesis was further supported by the significant increase of both photoactivation and photobleaching kinetics upon increase of local concentration through the variation of the probe/lipid ratio in the LUVs (1/100 to 1/1000) (Figure 4E). Indeed, the monitoring of the integrated signal over time combined to a kinetic model (see SI) allowed us to determine two kinetic constants k_Pa_ and k_Bl_ that were respectively assigned to the photoactivation and the photobleaching rates and yielded respectively the quantum yield of photoactivation ϕ_Pa_ and of photobleaching ϕ_Bl_ (Figure 4F). Interestingly, ϕ_Pa_ was found to be 7 to 10-fold higher than ϕ_Bl_ (Figure 4F). Overall, these experiments showed that Rhod-PM acts as a mild photosensitizer, able to oxidize LRhod-PM in an efficient manner when concentrated on a plasma membrane model through a self-triggered photooxidation cascade.

### Photoactivation of LRhod-PM in cell imaging

Once the proof of concept established, we assessed the self-triggered photooxidation cascade mechanism on cells using LRhod-PM. As controls, the plasma membrane staining of the probes were checked in HeLa cells using laser scanning confocal microscopy. While LRhod-PM did not produce any signal, Rhod-PM displayed a fast and wash-free fluorescent plasma membrane staining with high signal-to-noise ratio of 29 and a Pearson’s colocalization coefficient of 0.80 with MemBright-Cy5.5 (Figure 5).

**Figure 5.**
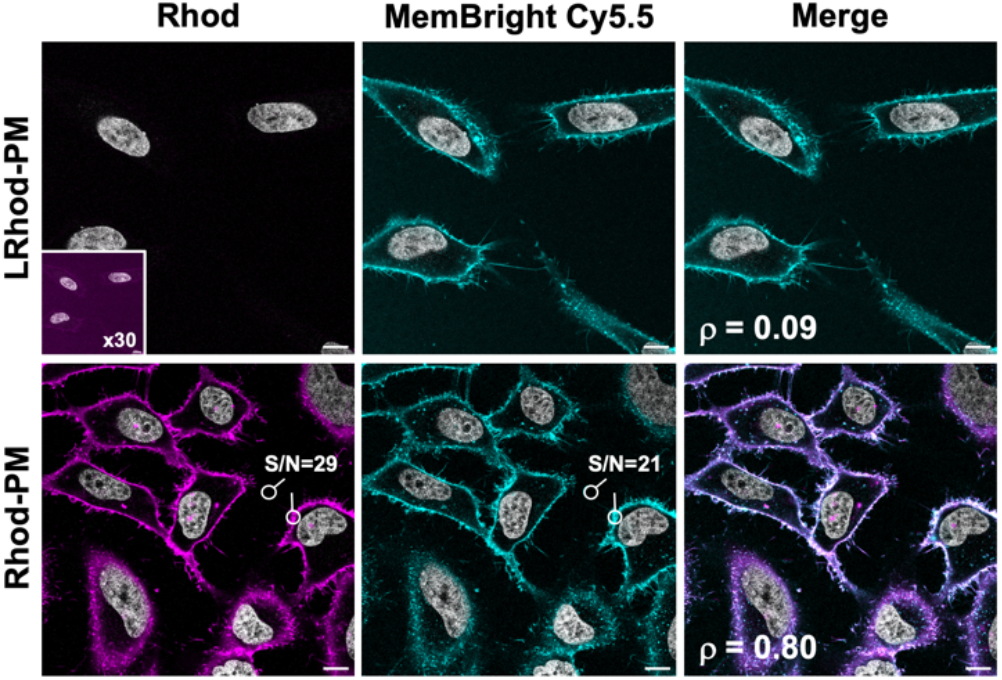
Plasma membrane staining with LRhod-PM and Rhod-PM. Laser scanning confocal microscopy images of live HeLa cells 5 min after addition of LRhod-PM or Rhod-PM (200 nM) in wash-free conditions. MemBright Cy5.5 was used as co-staining marker (200 nM). Nuclear staining (gray color) was done with Hoechst (5 µg/mL). Scale bar: 10 µm. ρ is the Pearson’s coefficient depicting the colocalization between the probes and MemBright Cy5.5. S/N is the signal to noise ratio.

Then, cells incubated with LRhod-PM were irradiated for 40 s at 552 nm on a zoomed region at higher laser power (10%) and the cells were imaged over time (Figure 6A). Upon irradiation, the fluorescence signal progressively increased (5-fold) to reach a plateau after ≈30 s (Figure 6B, Supplementary movie 1). The obtained image displayed a good signal-to-noise ratio of 7 and indicated an efficient plasma membrane localization as proven by the high Pearson’s colocalization coefficient with MemBright (Figure 6C).

**Figure 6.**
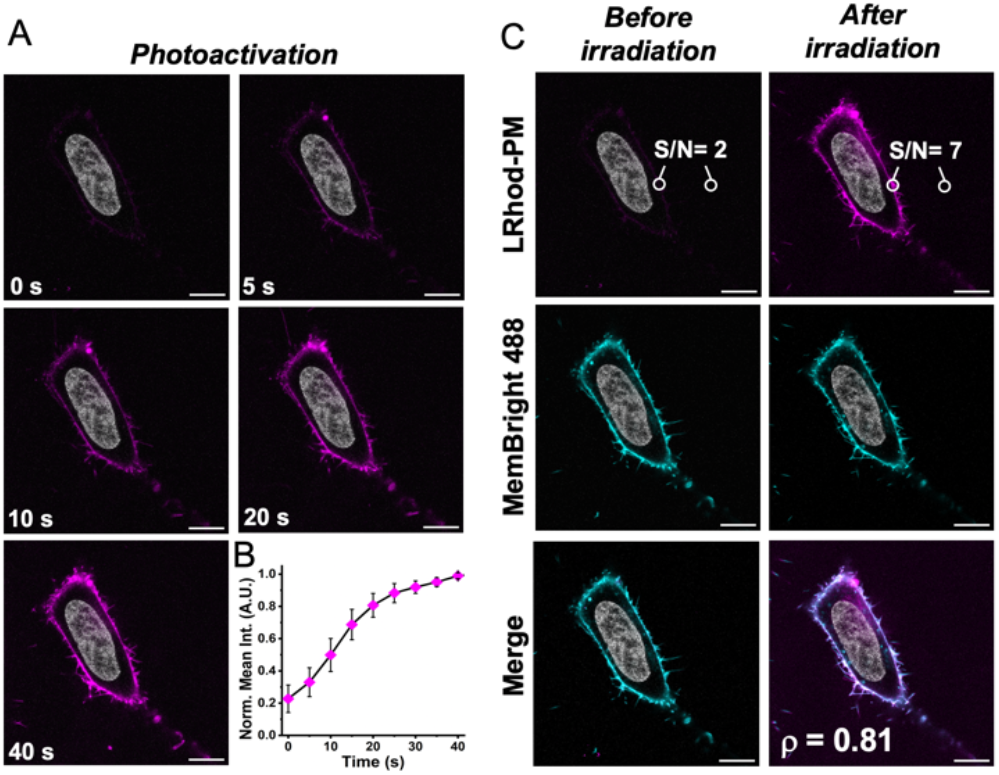
Photoactivation of LRhod-PM in HeLa cell. (A) Photoactivation of LRhod-PM (200 nM) upon 552 nm irradiation of a single cell (B) Increase of the mean fluorescence intensity upon irradiation. Fluorescence signal increase was measured on three different cells. (C) Laser scanning confocal microscopy images of HeLa cells with LRhod-PM (200 nM) and MemBright 488 (200 nM) before and after irradiation. Nuclear staining (gray color) was done with Hoechst (5 µg/mL). Scale bars: 10 µm. S/N is the signal to noise ratio.

Next, we confirmed that at the working concentration of 200 nM, both Rhod-PM and L-Rhod-PM were not cytotoxic nor photocytotoxic under irradiation using MTT cell viability assays in HeLa cells before and after irradiation (Figure S3). Overall, these cellular experiments successfully showed that L-Rhod can be used as an efficient photoactivatable plasma membrane probe without detrimental cytotoxic or photocytotoxic effects.

### Self-Triggered Photooxidation Cascade enables Live SMLM

Common PALM probes typically require UV photoactivation,^[10], [11], [14], [54]^ which is not preferable for live-cell applications. Conversely LRhod-PM possesses appealing feature for live-SMLM as it only requires a single laser in the visible range. Also, we assumed that the presented mechanism would find advantages in super-resolution imaging as it gathers the ideal conditions to perform live single-molecule localization microscopy (SMLM) in PALM conditions. Indeed, at the single-molecule level, an efficient photoactivation combined with an accelerated photobleaching would provide an efficient blinking profile and enable to reach a single-molecule regime where single emitters are temporally and spatially well separated.^[55]^ However, in live super-resolution imaging based on SMLM acquisition time should be low, *i*.*e*. the number of blinks should be relatively high to have an efficient reconstruction and to palliate the potential movement of the cell during imaging. Hela cells were then stained with LRhod-PM at the low concentration of 20 nM. In widefield microscopy, the low signal-to-noise ratio virtually did not enable the discrimination of the plasma membrane. Consequently, the cells and focal plan were found using a nuclear staining in an independent channel (Hoechst, λ_Ex_: 405 nm).

Upon irradiation of a region of interest (Figure 7A) at 532 nm (0.25 kW.cm^-2^) a high frequency of blinking event has been obtained and the single-molecule regime could be reached (See Supplementary movie 2). Importantly, the laser power density used here was significantly lower than many conventional SMLM experiments, typically 1 to 3 kW.cm^-2^.^[56]^ This blinking efficiency allowed to reconstruct the plasma membrane from a 120 s acquisition only (9k frames at 14 ms frame rate) and with a high gain of resolution (Figure 7B). Indeed, while the widefield image after irradiation provided a membrane thickness of 681 nm, the SMLM image indicated 77 nm (Figure 7C). Due to the high brightness of Rhod-PM and its efficient excitation with the 532 nm laser line, the blinking event displayed a rather high mean localization precision of 21.1 ± 4.6 nm (Figure 7D). Moreover, the localized photooxidation of LRhod provided a fairly stable number of localizations along with a constant number of photons over the time (Figure 7E and F).

**Figure 7.**
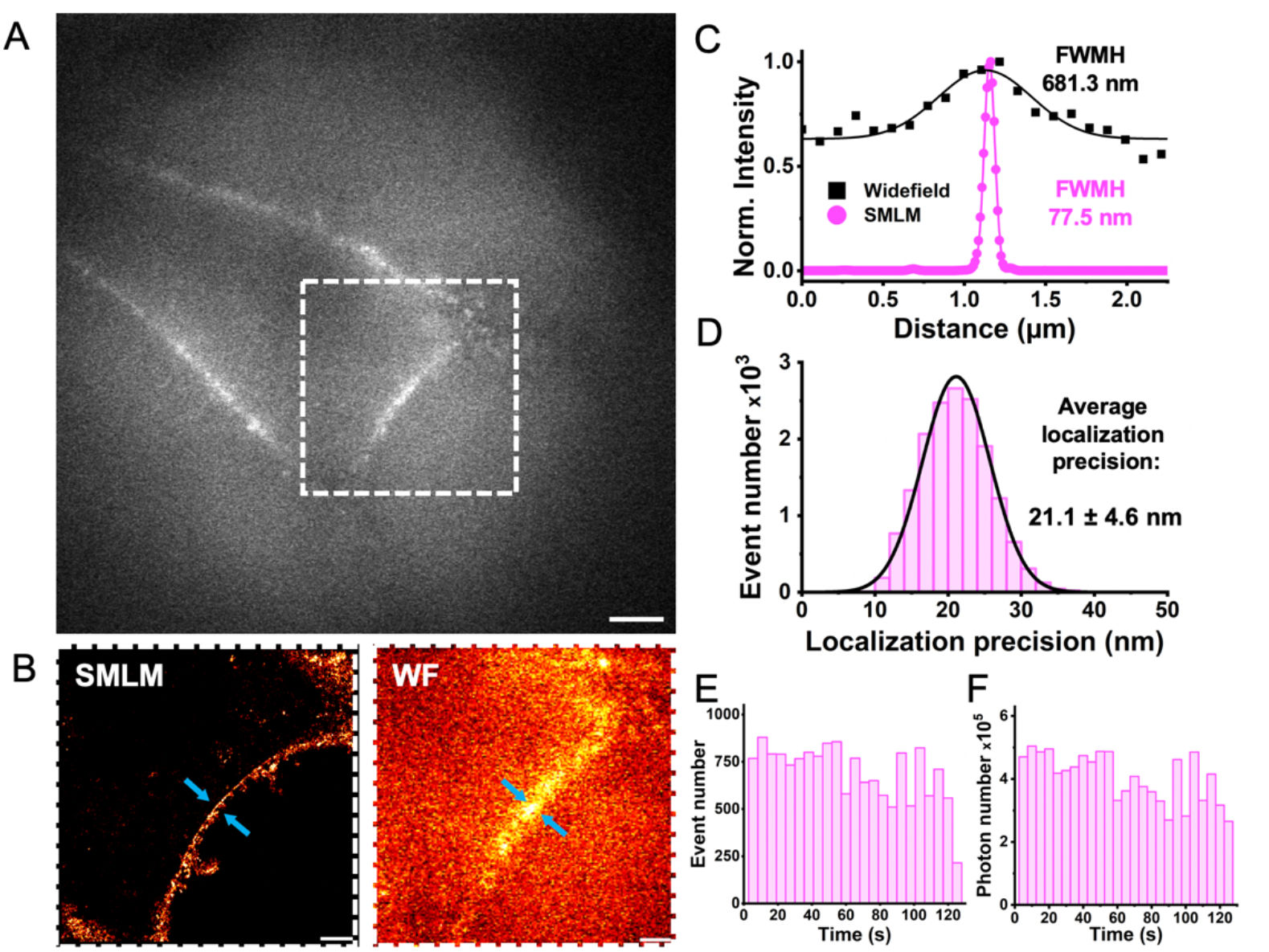
Live-cell SMLM with LRhod-PM (20 nM). (A) Widefield image after photoactivation. Scale bar: 5 µm. (B) Super-resolved image based on 16k localizations (left) and widefield image (right) of the region of interest. Scale bar: 2 µm. (C) Gaussian fits of intensity profiles of the position indicated by the arrow heads in the WF image and the corresponding SMLM image. (D) Average localization precision of detected events. (E) Detected event number per frame (F) Photon number over time. Binning: 5 secs.

In parallel, Rhod-PM was also evaluated for live SMLM. As expected, the same mean photon number per blink were obtained when using LRhod-PM and Rhod-PM since the former arises from the latter upon activation (Figure 8A). Although reconstruction after acquisition provided an image of the plasma membrane (Figure S4), the use of Rhod-PM evidenced some drawbacks compared to LRhod-PM. Despite the use of poly-lysine-coated glass, Rhod-PM, as a cationic probe is prone to stick on the glass surface which inducts a high background noise. This significantly reduces the localization density and the reconstructed image quality, which is depicted by the signal-to-noise ratio in widefield after irradiation (Figure 8B). Finally, while a cell stained with LRhod-PM reaches a single-molecule regime after only a few seconds, Rhod-PM, which is intrinsically fluorescent, needs to be photobleached for almost one minute, which thus considerably reduced the reservoir of available blinking probes and increases the phototoxicity. This was illustrated by the mean event number of LRhod-PM which is twice higher than those of Rhod-PM over the acquisitions (Figure 8C, Supplementary movie 2). Overall, LRhod-PM enabled live SMLM imaging of plasma membrane in HeLa cells via Self-Triggered Photooxidation Cascade mechanism.

**Figure 8.**
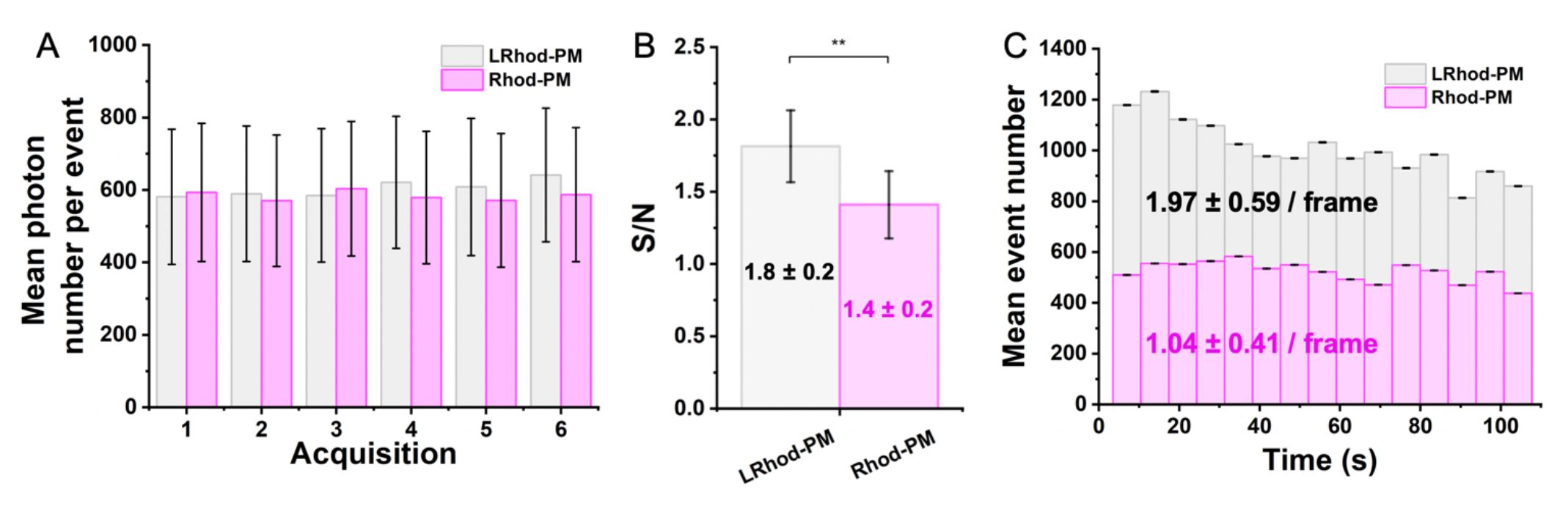
Live-cell SMLM performance of LRhod-PM and Rhod-PM (20 nM). Comparison were done on six different cells. (A) Mean photon number per event over 6 acquisitions. Mean event number per frame. Binning size: 7 sec (B) Signal-to-noise ratio in widefield after SMLM acquisition. **: p≤0.01 (C) Mean event number per frame.

## Conclusion

In conclusion, we herein synthesized LRhod-PM, a leuco-rhodamine plasma membrane-targeted probe which can be photoactivated and excited with a single laser upon irradiation in the visible range and can be used in live SMLM (PALM) to rapidly obtain images of membrane at the nanoscale. We first showed that the rhodamine Rhod-PM can act as a mild photosensitizer with a quantum yield of ^1^O_2_ generation lower than 1%. We then showed that ^1^O_2_ efficiently oxidizes the reduced form of rhodamine, called leuco-rhodamine. Then we demonstrated that LRhod-PM could get photoactivated by Rhod-PM in a concentration dependent manner. Similarly, the photobleaching of Rhod-PM is accelerated at high local concentration when embedded in lipid bilayer. This phenomenon, where a small number of Rhod-PM is sufficient to simultaneously trigger the photoactivation of L-Rhod and the photobleaching of Rhod-PM in an autocatalytic manner was called Self-Triggered Photooxidation Cascade (STPC). This mechanism, which was shown to be non-photocytotoxic to cells, was successfully applied to photoactivate the cell membrane in living cells as well as to perform live SMLM (PALM) imaging allowing to image the plasma membrane in 2 min with a high gain of resolution. This work shows the application of leuco-rhodamines in live SMLM and provides a new PM probe adapted to the orange channel using a single common laser line (530-560 nm) with low laser power and without any cytotoxic UV photoactivation. We expect that this mechanism will pave the way to a new generation of targeted and multicolor fluorescent probes adapted to challenging super-resolution imaging of live samples.

## Materials and methods

### Synthesis

All starting materials for synthesis were purchased from Merck (Darmstadt, Germany), BLD Pharm (Reinbeck, Germany) or TCI Europe (Zwijndrecht, Belgium) and used as received unless stated otherwise. NMR spectra were recorded on a Bruker Avance III 400 MHz or 500 MHz spectrometers. Mass spectra were obtained using an Agilent Q-TOF 6520 mass spectrometer. Protocols and characterization of all new compounds are described in the Supporting Information.

### Spectroscopy

Absorption and emission spectra were recorded at 20° C, 1 µM in MeOH on a UV-2700 spectrophotometer (Shimadzu) and a FluoroMax-4 spectrofluorometer (Horiba Jobin Yvon) equipped with a thermo-stated cell compartment. For standard recording of fluorescence spectra, the emission was collected 5 nm after the excitation wavelength. All the spectra were corrected from the wavelength-dependent response of the detector. The fluorescence quantum yields ϕ_*F*_ was determined by comparison with rhodamine B in water,^[57]^ as reference according to its excitation and emission wavelengths according to the following equation:

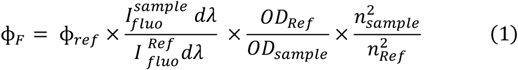

with ***I***_***fluo***_ : integration of the fluorescence signal, ***OD*** : optical density at the excitation wavelength, and ***n*** : refraction index of the solvent.

### Laser spectroscopy

Laser spectroscopy studies were performed using 3 × 3 mm optical path length quartz cuvettes of 45 µL. Excitation was provided by a continuous wave laser diode (405, 532 or 638 nm, Oxxius, Lannion, France) and photons were detected by a QE pro spectrometer (Ocean Optics, USA). All measurements were performed at room temperature.

### Singlet oxygen quantum yield

5 µM of photosensitizer (Rhod-DA, Rhod-PM or rose bengal) was added to a freshly prepared solution of 100 µM chemical acceptor (DPBF) in MeOH. The mixture was irradiated using 532 nm laser (43 mW.cm^-2^). DPBF fluorescence intensity was recorded over time thanks to 405 nm laser (0.4 mW.cm^-2^). The emission spectra were acquired to obtain the decreasing slope of DPBF fluorescence intensity. The experiment was repeated twice. For correct linear fitting, only the first points were used, keeping DPBF consumption below 20%. The consumption of DPBF can be expressed as^[58]^:

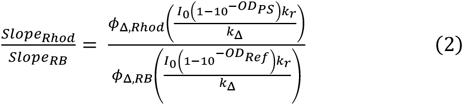

with *slope* : linear slope of the integrated fluorescence intensity disappearence of DPBF thanks to photosensitizer (PS) or reference (Ref), *ϕ*_Δ_ : singlet oxygen quantum yield, *I*_0_ : irradiance of the light source, (1 ™ 10^™OD^) : absorption factor of the sample, *k*_*r*_ : reaction with the chemical acceptor, *k*_Δ_ : solvent-mediated deactivation.

As chemical acceptor concentration and light irradiance are equal for the sample (Rhod-DA or Rhod-PM) and reference (rose bengal, ϕ_Δ,MeOH_= 0.77^[49]^), the singlet oxygen quantum yield of rhodamine dyes can be expressed as:

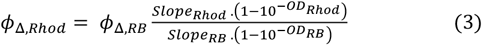

### Exogenous singlet oxygen generation

LRhod-DA (1 µM) and methylene blue (1 µM, ϕ_Δ,MeOH_= 0.50^[49]^) were irradiated for 30 minutes using 638 nm laser (90 mW.cm^-2^). The emission spectra of the oxidized form Rhod-DA were acquired at t0 and t30 minutes upon brief 532 nm laser excitation (68 mW.cm^-2^).

### Lipid vesicles

LUVs were obtained by the extrusion method as previously described.^[59]^ Briefly, a suspension of multilamellar vesicles was extruded by using a Lipex Biomembranes extruder (Vancouver, Canada). The pore size of the filters was first 0.2 μm (10 passages) and thereafter 0.1 μm (10 passages) to generate monodisperse LUVs with a mean diameter of 0.1 μm as measured with a Malvern Zetamaster 300 (Malvern, U.K.). LUVs were composed of DOPC for a final concentration of 200 µM. For the experiment with LUVs, the probe:lipid ratio was set to 1:100, 1:250, 1:500 and 1:1000.

### Photooxidation studies in lipid vesicles. Determination of kinetic constants and photooxidation quantum yields

LRhod-PM was embedded into LUVs at different molar ratios by mixing LRhod-PM (200 nM) and lipid vesicles (lipid concentration: 20, 50, 100 or 200 µM) for 30 min at room temperature. The mixture was irradiated using 532 nm laser (72 mW.cm^-2^) and the fluorescence intensity was recorded over time. Each condition was recorded in triplicate. Kinetic rate constants of photoactivation and photobleaching processes were determined by fitting the integrated fluorescence signal IF of Rhod-PM over time according to the equation (4).^[60]^

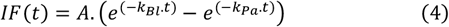

with *k*_*Bl*_ : constant rate of photobleaching and *k*_*pa*_: constant rate of photoactivation.

The quantum yields of photomodulation Φ_Pa_ and Φ_Bl_ were calculated from the rate constants k_Pa_ and k_Bl_ respectively according to the equation (5):

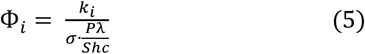

with P: laser power used for irradiation (W), λ: wavelength of irradiation (m), S: irradiated surface (here S= 0.15 cm^2^), h: Planck’s constant, c: lightspeed (m.s^-1^) and σ: absorption cross section (cm^2^). The latter is defined by the following equation:

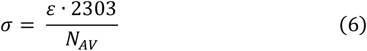

with ε: molar-absorption coefficient of the dye at the excitation wavelength (L.mol^-1^.cm^-1^) and the Avogadro’s constant N_AV_ (mol_-1_). More information is provided in Supplementary.

### Photooxidation studies in HPLC. Determination of chemical yield

LRhod-DA after irradiation using 532 nm laser (71 mW.cm^-2^) was subjected to a reversed-phase HPLC system with a Uptisphere 120Å C18-ODB column and a Waters 2996 photodiode array detector. Analysis was done using a gradient elution starting from 100% H_2_O (+ 0.1% TFA) to 100% acetonitrile in 30 min. The signal of LRhod-DA and Rhod-DA were monitored at 254 nm. Subsequently, the quantification of Rhod-DA was calculated from the peak area correlated to the calibration curve presented in Figure S2. The calibration was made with the analysis of 5 different concentrations in triplicate (0.1, 0.5, 1, 5 and 10 µM). Each triplicate results from an independent weighting of Rhod-DA.

### Cytotoxicity and Photocytoxicity

Cytotoxicity was evaluated by MTT assay (3-(4,5-dimethylthiazol-2-yl)-2,5-diphenyltetrazolium bromide).^[61]^ A total of 2.10^4^ HeLa cells/well were seeded in a 96-well plate 24 h prior to the experiment in Minimum Essential medium (MEM, Gibco Lifetechnologies) complemented with 10% FBS, Penicillin-Streptomycin (100 µg/mL), L-Glutamine (2 mM), non-essential amino acids (1 mM), sodium pyruvate (1 mM) and were incubated in a 5% CO_2_ incubator at 37°C. After medium removal, an amount of 100 µL growing media containing 200 nM of Rhod-PM or LRhod-PM were added to the HeLa cells and incubated for 30 min at 37°C (5% CO_2_). As controls, the cells were incubated with growing media (+ 0.1% DMSO) as positive control and Triton™ X-100 (1%) at negative control of cell viability. After 30 min of dye incubation, the medium was replaced by 100 µL of growing media containing 10 % MTT solution in PBS (4 mg.mL^-1^) and the cells were incubated for 3 h at 37°C (5% CO_2_). MTT solution was then removed, replaced by 100 µL of DMSO and gently shaken for 10 sec at room temperature. The absorbance at 570 nm was recorded. Each condition was tested in octuplicate, the percentage of cell viability was calculated compared to the control positive control. The photocytotoxicity was evaluated with the same procedure as cytotoxycity, differing by an irradiation step of the cells with 525 nm Green LED Array Light Source (Thorlabs, 0.3 mW) for 30 min after adding the probe instead of incubating in the dark.

### Cell sample preparation

HeLa cells were incubated in Minimum Essential medium (MEM, Gibco Lifetechnologies) complemented with 10% FBS, Penicillin-Streptomycin (100 µg/mL), L-Glutamine (2 mM), non-essential amino acids (1 mM), sodium pyruvate (1 mM) and were incubated in a 5% CO_2_ incubator at 37°C. For the imaging experiments, cells were seeded onto a 35 mm IBiDi live cell plates at a density of 90 × 10^3^ cells/well 24 h before the microscopy measurements.

### Colocalization

For a nuclear staining, the medium was replaced by Hoechst 33342 (Thermo Fisher, 5 µg/mL) in Opti-MEM (Gibco-Invitrogen) and the cells were incubated for 30 minutes at 37°C. Cells were then incubated with 200 nM of plasma membrane probe in Opti-MEM for 5 minutes (LRhod-PM, Rhod-PM or MemBright Cy5.5 as a co-staining marker). Without any washing step, cells were imaged with a Leica TSC SP8 laser scanning confocal microscope with a 63x/1.40 oil objective. The microscope settings were: 405 nm laser (5%) for excitation of Hoechst 33342, emission was collected between 410 and 470 nm; 552 nm laser (1%) for excitation of Rhod-PM and LRhod-PM, emission was collected between 560 and 630 nm; 638 nm laser (1%) for excitation of MemBright Cy5.5, emission was collected between 650 and 800 nm. The Pearson’s coefficients were obtained using the imageJ plugin: Colocalization Finder.

### Photoactivation in cells

For a nuclear staining, the medium was replaced by Hoechst 33342 (Thermo Fisher, 5 µg/mL) in Opti-MEM (Gibco-Invitrogen) and the cells were incubated for 30 minutes at 37°C. Cells were then incubated with 200 nM of plasma membrane probe in Opti-MEM for 5 minutes (LRhod-PM or MemBright 488 as a co-staining marker). Without any washing step, cells were imaged with a Leica TSC SP8 laser scanning confocal microscope with a 63x/1.40 oil objective. The microscope settings were: 405 nm laser (5%) for excitation of Hoechst 33342, emission was collected between 410 and 470 nm; 488 nm laser (1%) for excitation of MemBright 488, emission was collected between 495 and 545 nm; 552 nm laser (1%) for excitation of MemBright Cy5.5, emission was collected between 560 and 650 nm. Cells were found with nuclear staining. The activation process was done at 10% 552 nm laser and was monitored through the evolution of the intensity histogram, the irradiation scans were stopped when the histogram presented no more evolution. The Pearson’s coefficients were obtained using the imageJ plugin: Colocalization Finder.

### SMLM imaging

Cells were seeded onto a 35 mm IBiDi polylysine coated plates at a density of 90.10^3^ cells/well 24 h before the microscopy measurements. All samples were imaged using a home-built set-up, based on a Nikon Eclipse Ti microscope with 100x 1.49 NA oil-immersion objective. All single-molecule images were aquired using 532 nm illumination (250 W.cm^-2^). An acousto-optic tunable filter (AOTF; Opto-Electronic) was used to modulate the laser power. The signal was detected with an EM-CDD camera (ImagEM, Hamamatsu) through a dichroic filter (Di02-532/635-t1-25x36, Semrock) and a 532 notch filter (NF03-532E-25, Semrock). Regions of interest (ROIs) were quickly identified thanks to nuclear staining (Hoechst 33342, 0.3 µg/mL) under 405 nm illumination. A 13.9 ms acquisition time per frame and 10,000 frames were collected for each movie. SMLM movies were analyzed using ThunderSTORM plug-in on ImageJ software (version 1.54i).^[62]^ Images were filtered with a wavelet filer (B-spline order 3, scale 2.0), approximate localization of the molecules was performed with a local maximum method (peak intensity threshold of 2*standard deviations, connectivity 8-neighbourhood), and sub-pixel localization of molecules was performed with a maximum likelihood fitting method (PSF integrated Gaussian method, fitting radius 3 px, initial sigma 1.6 px). Post-processing was performed by localizations filtering: 20 < σ < 200 and 200 < photon number < 1000. Reconstructed images were produced using a normalized Gaussian method with a lateral uncertainty of 20 nm. Intensity profiles were fitted with Gaussian fit to determine FWMH values.

## Supporting information

Movie1_photoactivation of L-Rhod-PM

Movie2_SMLM acquisition Rhod-PM vs LRhod-PM

## Supporting Information

Protocols of synthesis and characterizations (^1^H NMR ^13^C NMR, HPLC-High-resolution mass spectrometry) of the synthesized probes can be found in supporting information as well as the method to determine kinetic constants, additional experiments and supplementary figures.

Supplementary movie1. Photoactivation of LRhod-PM

Supplementary movie2. SMLM acquisition Rhod-PM vs LRhod-PM

## Acknowledgements

This work was founded by the the Agence Nationale de la Recherche: ANR 5D-SURE ANR-21-CE42-0015. The authors would like to thank Pr. Pascal Didier, Dr. Andrey S. Klymchenko and Dr. Alexandre Specht for their help.

## Note

Kyong Fam current address is : Department of Immunology and Microbiology, Scripps Research, La Jolla, California 92037, United States

## Entry for the Table of Contents

**Synthesis, protocol:**

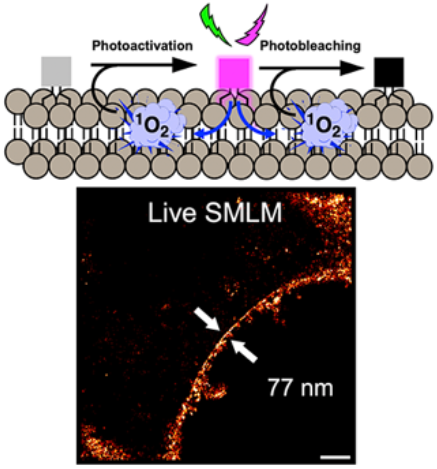

Herein we establish a new photoswitching concept called Self-Triggered Photooxidation Cascade (STPC). Rhodamine dyes have mild photosensitizing properties that enable subsequent photoactivation and photobleaching of a leuco-rhodamine plasma membrane probe. Upon laser irradiation in the visible range, we successfully performed live SMLM (Single-Molecule Localization Microscopy), allowing to image the plasma membrane at the nanoscale.

Institute and/or researcher Twitter usernames: @sonia_pfister, @vleberruyer, @KyongFAM, @Mayeul_Collot

## Supplementary Information

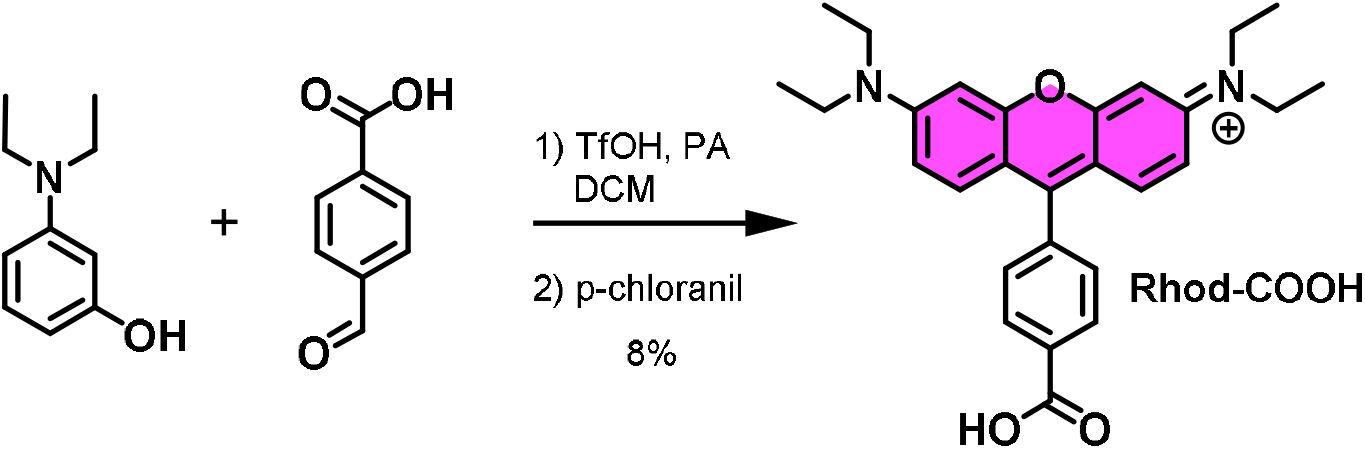

4-carboxybenzaldehyde (300 mg, 2 mmol, 1 eq.) and diethylaniline (826 mg, 5 mmol, 2.5 eq.) were dissolved in DCM (5 mL) and propionic acid (2 mL). Triflic acid (53 µL, 0.6 mmol, 0.3 eq.) was added and reaction mixture was stirred at room temperature for 24 h. As condensated product was formed and aldehyde completely consumed, chloranil (491 mg, 2 mmol, 1 eq.) was added and reaction stirred for further 24 h at room temperature. Solvents were evaporated and product was purified by flash chromatography using solid deposite (DCM/MeOH 98/2 to 90/10) to obtain **Rhod-COOH** as a pink solid with metallic sheen (72 mg, 8%).

^1^H NMR (400 MHz, MeOD) δ 8.28 (d, *J* = 8.3 Hz, 2H, 2H_ar, benzene_), 7.54 (d, *J* = 8.3 Hz, 2H, 2H_ar, benzene_), 7.30 (d, *J* = 9.5 Hz, 2H, 2H_ar, xanthene_), 7.05 (dd, *J* = 9.5, 2.4 Hz, 2H, 2 H_ar, xanthene_), 6.95 (d, *J* = 2.4 Hz, 2H_ar, xanthene_), 3.67 (q, *J* = 7.1 Hz, 8H, 4xN-C<u>H_2_</u>), 1.30 (t, *J* = 7.1 Hz, 12H ; 4xN-CH_2_-C<u>H_3_</u>)

^13^C NMR (500 MHz, MeOD) δ 169.36 (<u>C</u>OOH), 159.44 (C_ar_), 157.68 (C_ar_), 157.19 (C_ar_), 137.64 (C_ar_), 134.54 (C_ar_), 132.79 (C_ar_), 131.06 (C_ar_), 130.90 (C_ar_), 115.62 (C_ar_), 114.25 (C_ar_), 97.47 (C_ar_), 46.88 (<u>C</u>H_2_-N), 12.82 (<u>C</u>H_3_-CH_2_-N).

HRMS (ESI^+^): *m/z* calculated for C_28_H_31_N_2_O_3_^+^ M^+^: 443.2335, found 443.2345.

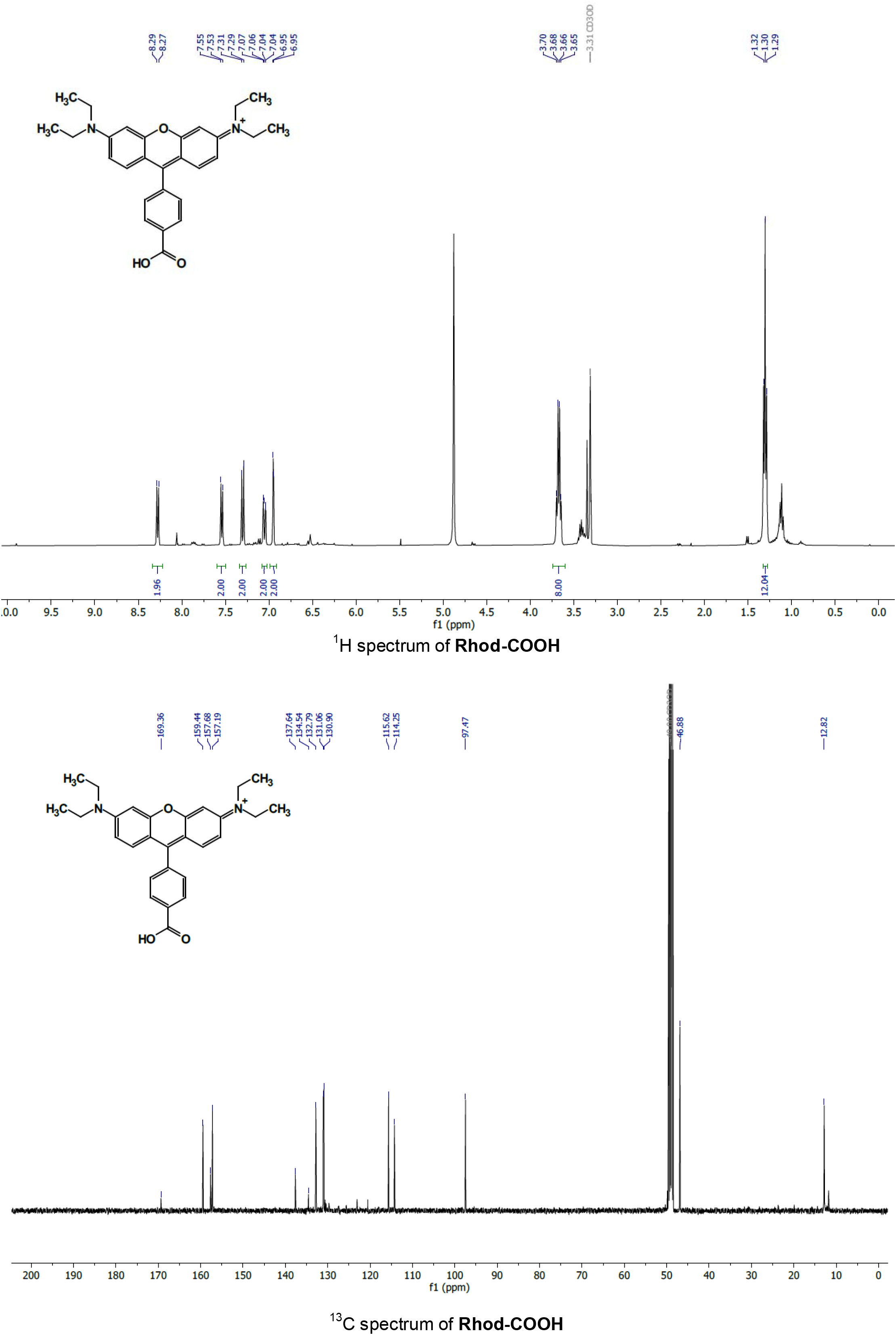
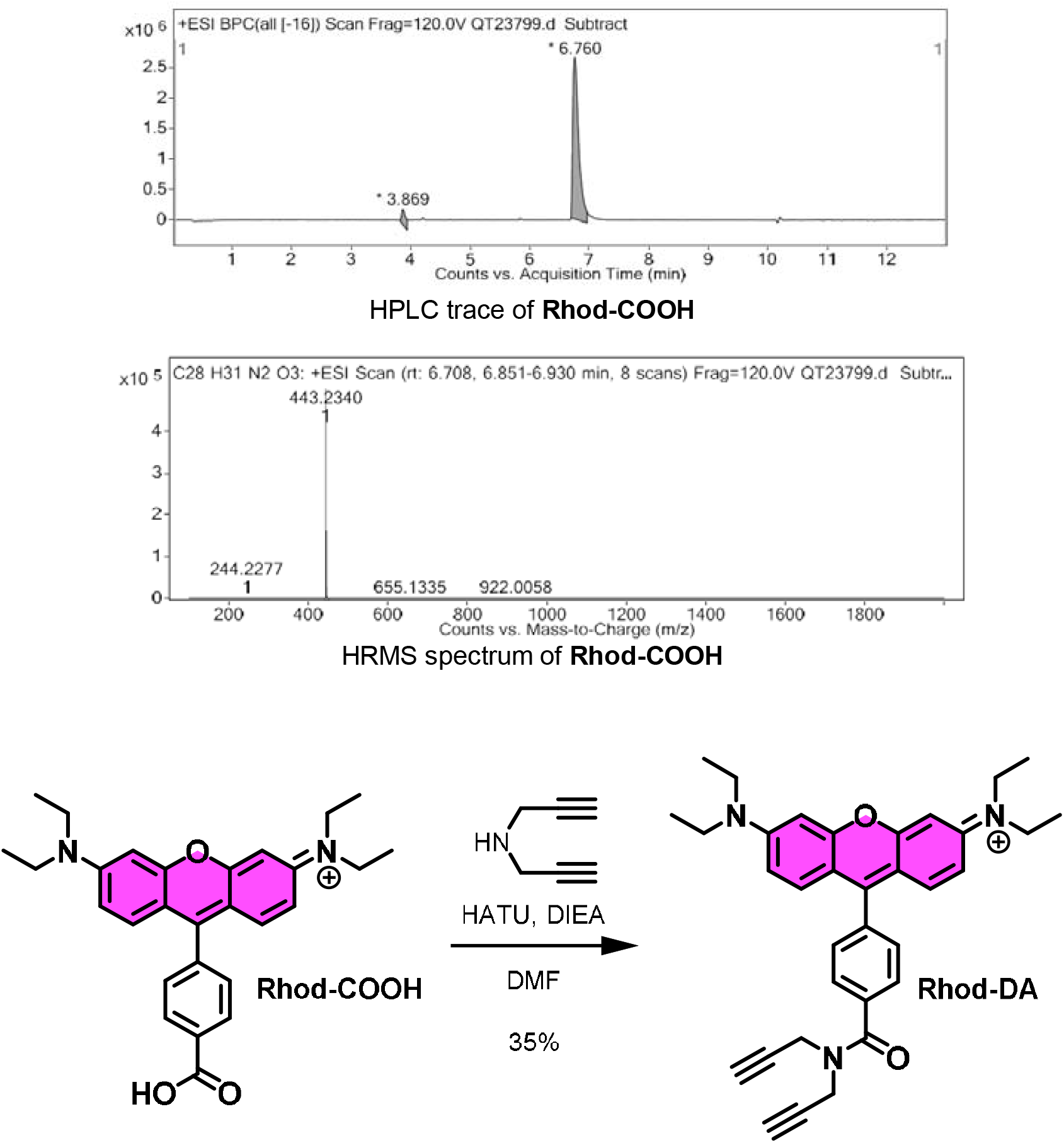

**Rhod-COOH** (72 mg, 0.16 mmol, 1 eq.) was dissolved in degassed DMF (5 mL). Dipropargylamine (20 µL, 0.2 mmol, 1.2 eq) was added followed by DIEA (85 μL, 0.5 mmol, 3 eq) and HATU (74 mg, 0.2 mmol, 1.2 eq). Reaction mixture was stirred at room temperature overnight. DMF was evaporated under vacuum. Product was extracted with DCM, washed with HCl 1M, sat. NaHCO_3_, brine and dried over MgSO4. DCM was removed under vacuum and product was purified by flash chromatography (DCM/MeOH 98/2 to 90/10) to obtain **Rhod-DA** as a pink solid with metallic sheen (29.4 mg, 35%).

^1^H NMR (400 MHz, MeOD) δ 7.82 (d, *J* = 8.3 Hz, 2H, 2H_ar, benzene_), 7.60 (d, *J* = 8.3 Hz, 2H, 2H_ar, benzene_), 7.37 (d, *J* = 9.5 Hz, 2H, 2H_ar, xanthene_), 7.08 (dd, *J* = 9.5, 2.5 Hz, 2H, 2H_ar, xanthene_), 6.98 (d, *J* = 2.5 Hz, 2H, 2H_ar, xanthene_), 4.55 – 4.30 (m, 4H, 2xN-C<u>H_2_</u>-CCH), 3.69 (q, *J* = 7.1 Hz, 8H, 4xN-C<u>H_2_</u>-CH_3_), 2.97 – 2.75 (m, 2H, 2xC≡C<u>H</u>), 1.31 (t, *J* = 7.1 Hz, 12H, 4xN-CH_2_-C<u>H_3_</u>).

^13^C NMR (400 MHz, MeOD) δ 171.91 (<u>C</u>OOH), 159.46 (C_ar_), 157.36 (C_ar_), 157.17 (C_ar_), 137.74 (C_ar_), 135.78 (C_ar_), 132.86 (C_ar_), 131.19 (C_ar_), 128.58 (C_ar_), 115.66 (C_ar_), 114.30 (C_ar_), 97.48 (<u>C</u>≡CH), 78.70 (C≡<u>C</u>H), 46.89 (<u>C</u>H_2_-N), 12.82 (<u>C</u>H_3_-CH_2_-N).

HRMS (ESI^+^): *m/z* calculated for C_34_H_35_N_3_O_2_^+^ [M+H]^+^: 518.2807, found 518.2817.

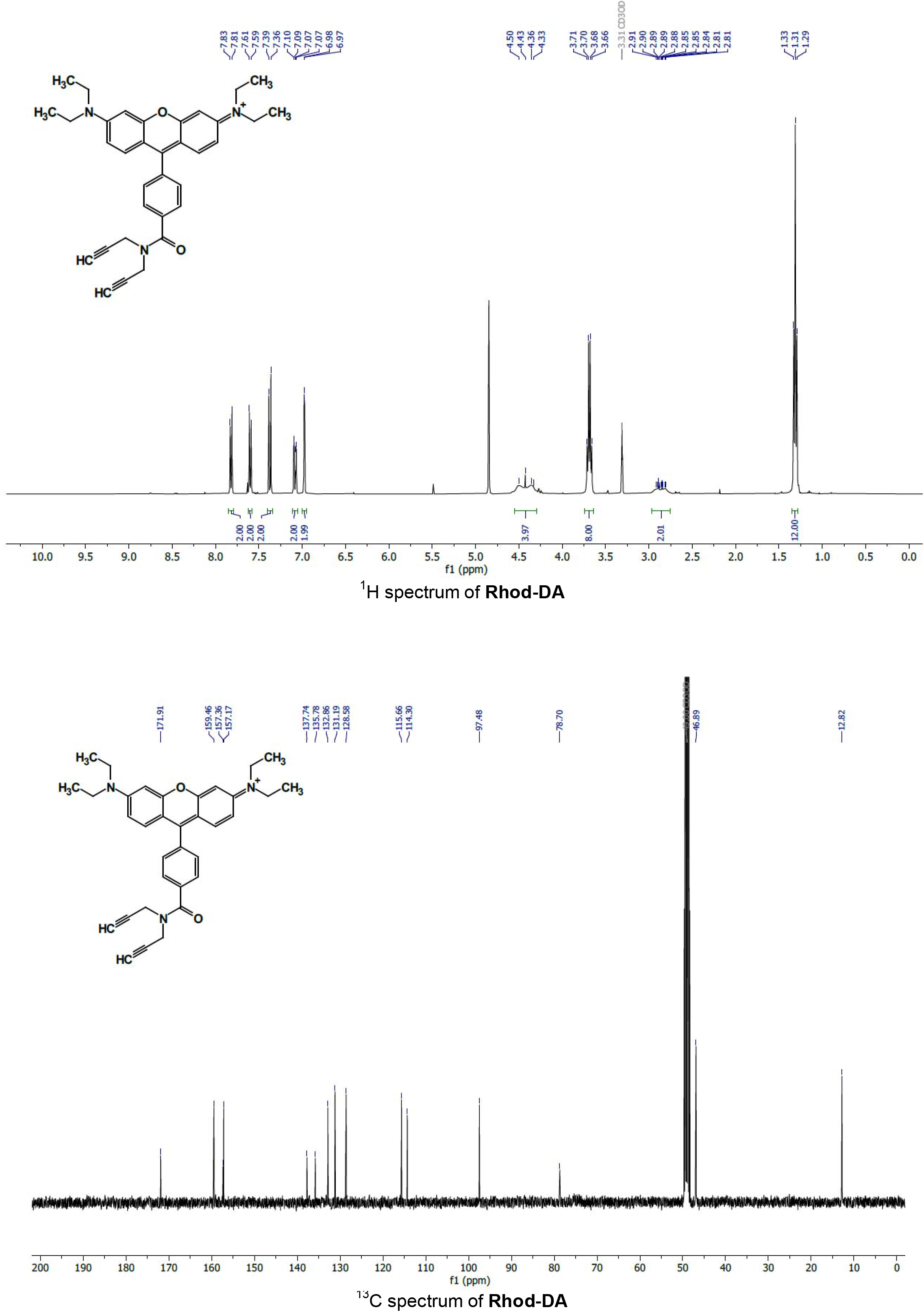
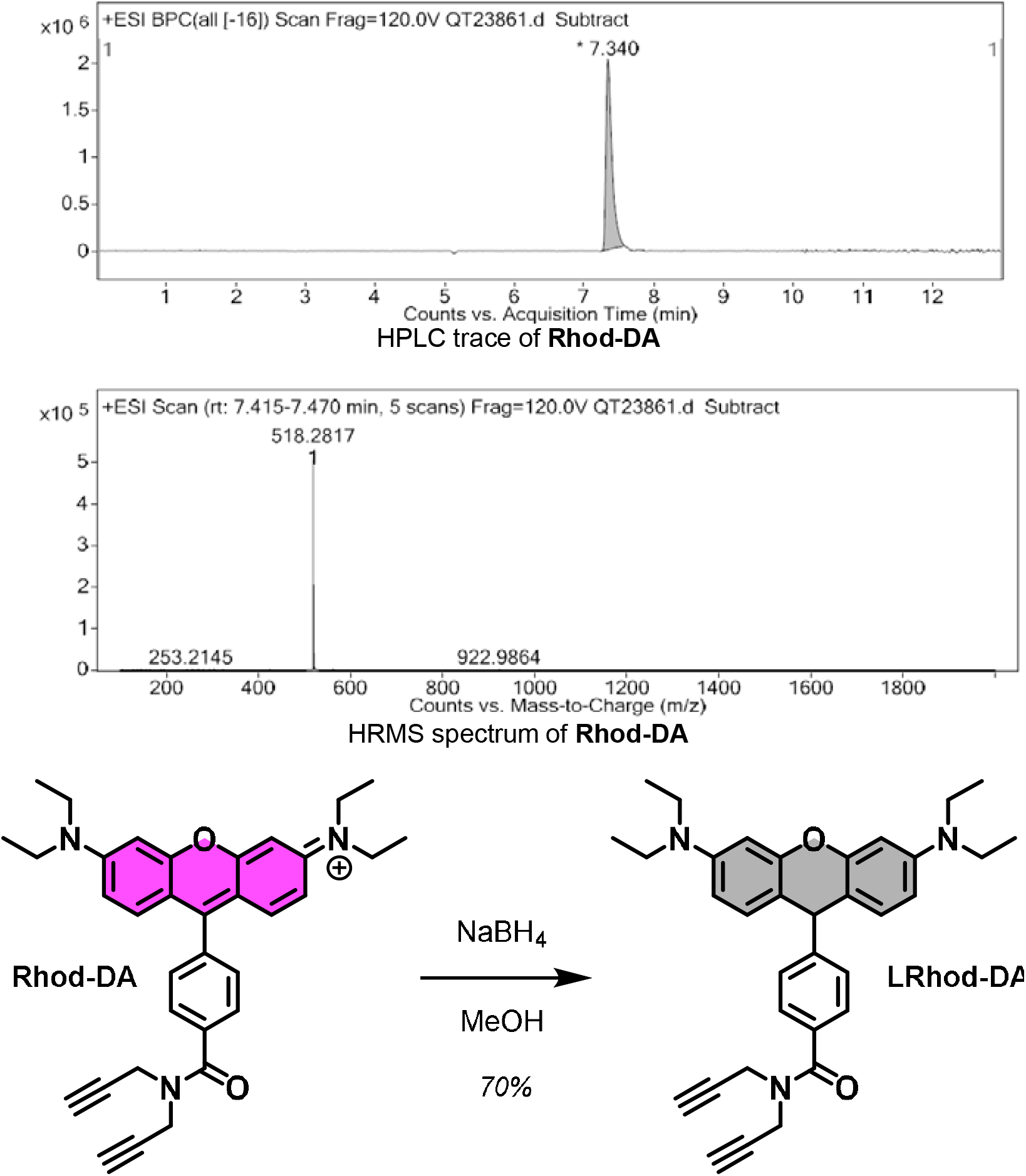

**Rhod-DA** (15 mg, 0.03 mmol, 1 eq.) was dissolved in anhydrous MeOH (2 mL) under argon atmosphere. NaBH_4_ (4.1 mg, 0.11 mmol, 4 eq.) was added and reaction mixture was stirred at room temperature for a few minutes and became light brown. HCl 1M was added until pH≈4 and product was extracted 3 times with EtOAc. Combined organic phases were dried over MgSO_4_ and concentrated under vacuum. Product was purified by flash chromatography (Heptane/EtOAc, 90/10 to 50/50) to obtain **LRhod-DA** as a light pink solid (9.9 mg, 70%). HRMS traces shows a small portion of oxidized **LRhod-PM** (**Rhod-PM**).

^1^H NMR (400 MHz, CDCl_3_) δ 7.47 (d, *J* = 8.2 Hz, 2H, 2H_ar, benzene_), 7.27 (d, *J* = 8.2 Hz, 2H, 2H_ar, benzene_), 6.78 (d, *J* = 8.6 Hz, 2H, 2H_ar, xanthene_), 6.38 (d, *J* = 2.6 Hz, 2H, 2H_ar, xanthene_), 6.32 (dd, *J* = 8.6, 2.6 Hz, 2H, 2H_ar, xanthene_), 5.08 (s, 1H, H_xanthene_), 4.51 – 4.13 (m, 4H, 2xN-C<u>H_2_</u>-CCH), 3.33 (q, *J* = 7.0 Hz, 8H, 4xN-C<u>H_2_</u>), 2.36 – 2.24 (m, 2H, 2xC≡C<u>H</u>), 1.16 (t, *J* = 7.0 Hz, 12H, 4xN-CH_2_-C<u>H_3_</u>).

^13^C NMR (500 MHz, CDCl_3_) δ 170.99 (<u>C</u>OOH), 152.24 (C_ar_), 151.05 (C_ar_), 147.97 (C_ar_), 132.22 (C_ar_), 130.44 (C_ar_), 128.80 (C_ar_), 127.82 (C_ar_), 111.29 (C_ar_), 107.59 (C_ar_), 98.86 (<u>C</u>≡CH), 78.41 (C≡<u>C</u>H), 44.54 (<u>C</u>H_2_-N), 42.87 (C_leuco_), 12.80 (<u>C</u>H_3_-CH_2_-N).

HRMS (ESI^+^): *m/z* calculated for C_34_H_37_N_3_O_2_ [M+H]^+^: 520.2964, found 520.2982.

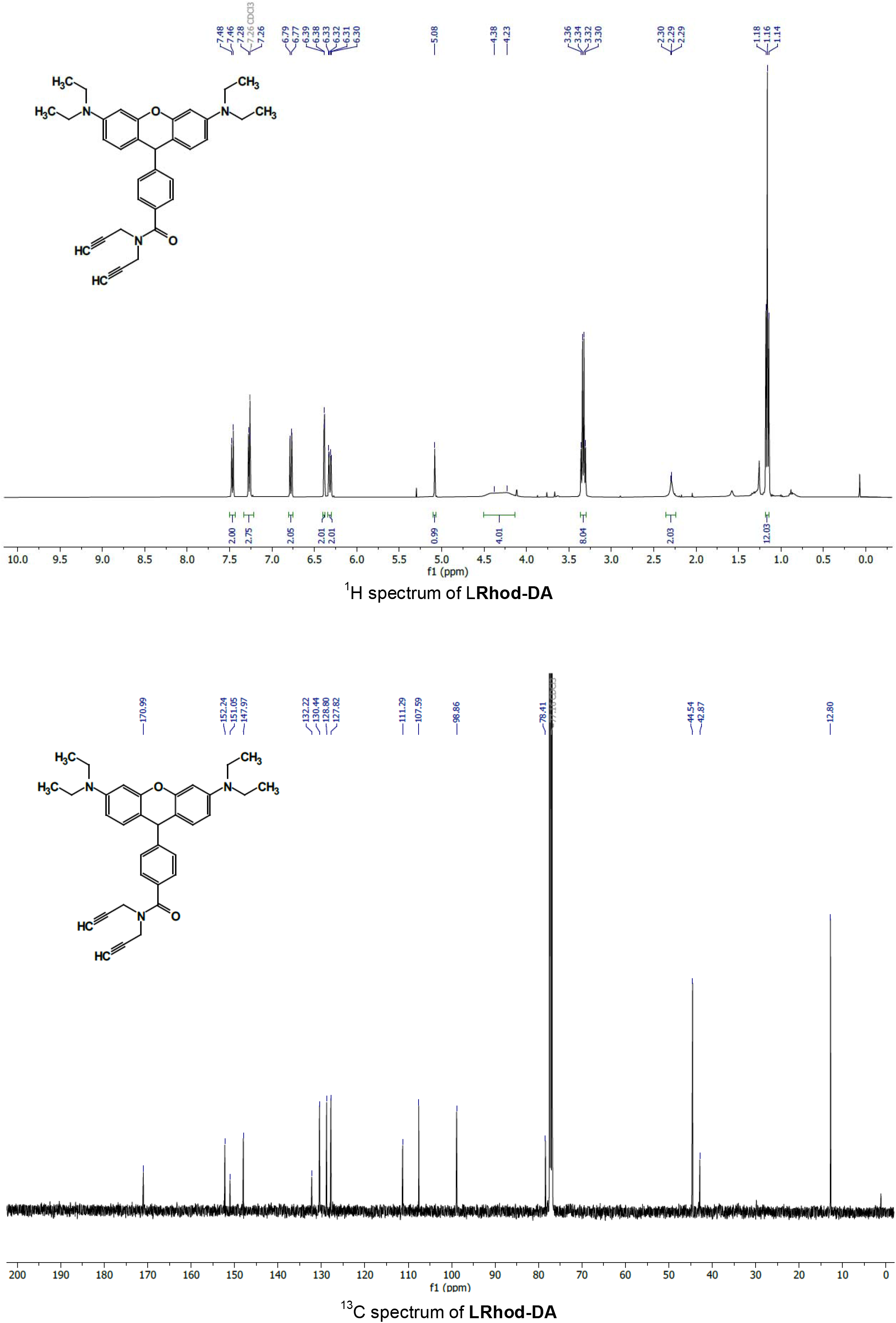
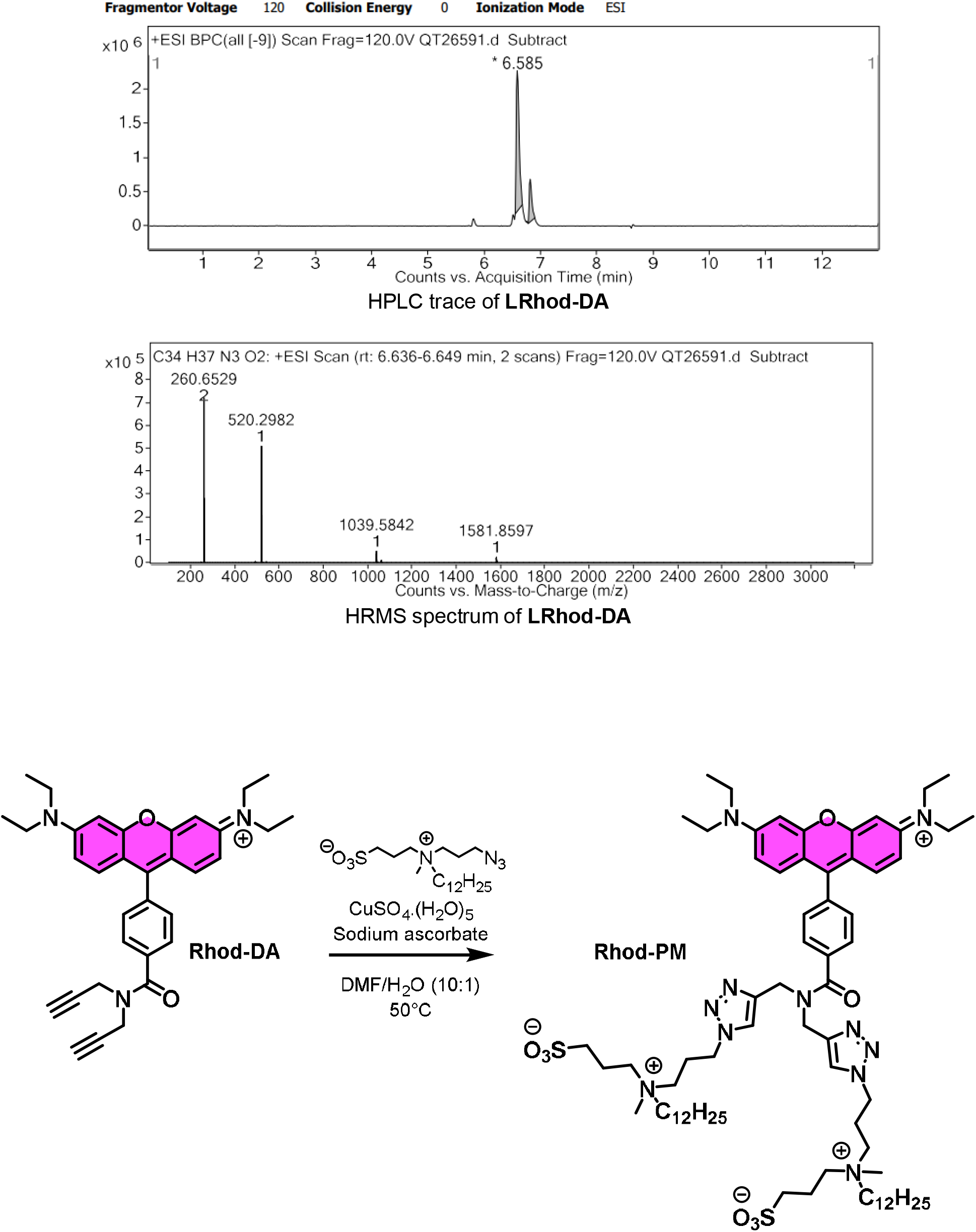

3-((3-azidopropyl)(dodecyl)(methyl)ammonio)propane-1-sulfonate was synthesized according to literature.^1^

**Rhod-DA** (25 mg, 0.048 mmol, 1 eq.) was dissolved in degassed DMF (2 mL). 3-((3-azidopropyl)(dodecyl)(methyl)ammonio)propane-1-sulfonate (48.4 mg, 0.116 mmol, 2.4 eq.) was added. Copper sulfate pentahydrate (5 mg) and sodium ascorbate (5 mg) were dissolved in an eppendorf with water (200 μL) and the solution was vortexed until the mixture turn yellow/orange. The content of the eppendorf was then added to the mixture and the reaction was stirred at 50°C for 1 h. Mixture was filtrated through a pad of celite and concentrated under vaccum. Crude product was purified by gel filtration chromatography Sephadex® LH-20 with DCM/MeOH (1/1) as eluent to obtain **Rhod-PM** as a pink solid with metallic sheen (63.4 mg, 95%).

HRMS (ESI^+^): *m/z* calculated for C_72_H_116_N_11_O_8_S_2_^+^ M^+^: 1326.8449, found 1326.8428.

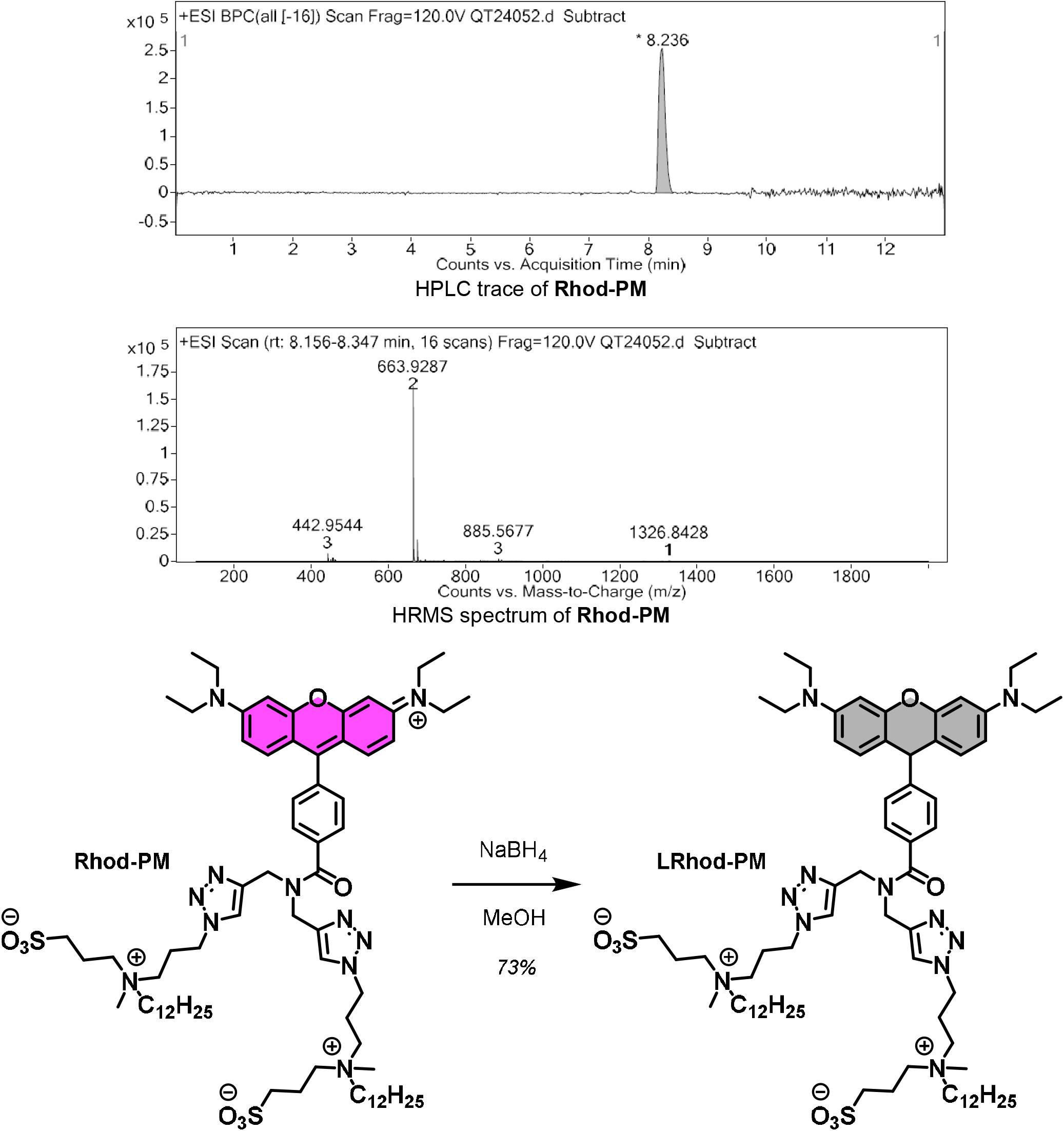

**Rhod-PM** (30 mg, 0.02 mmol, 1 eq.) was dissolved in anhydrous MeOH (5 mL) under argon atmosphere. NaBH_4_ (3.4 mg, 0.09 mmol, 4 eq.) was added and reaction mixture was stirred at room temperature for a few minutes and became light brown. Crude product was purified by gel filtration chromatography Sephadex® LH-20 with DCM/MeOH (1/1) as eluent to obtain **LRhod-PM** as a light pink solid (22 mg, 73%). HRMS trace shows a small portion of oxidized **LRhod-PM** (**Rhod-PM**).

HRMS (ESI^+^): *m/z* calculated for C_72_H_117_N_11_O_8_S_2_ [M+H]^+^: 1328.8606, found 1328.8531.

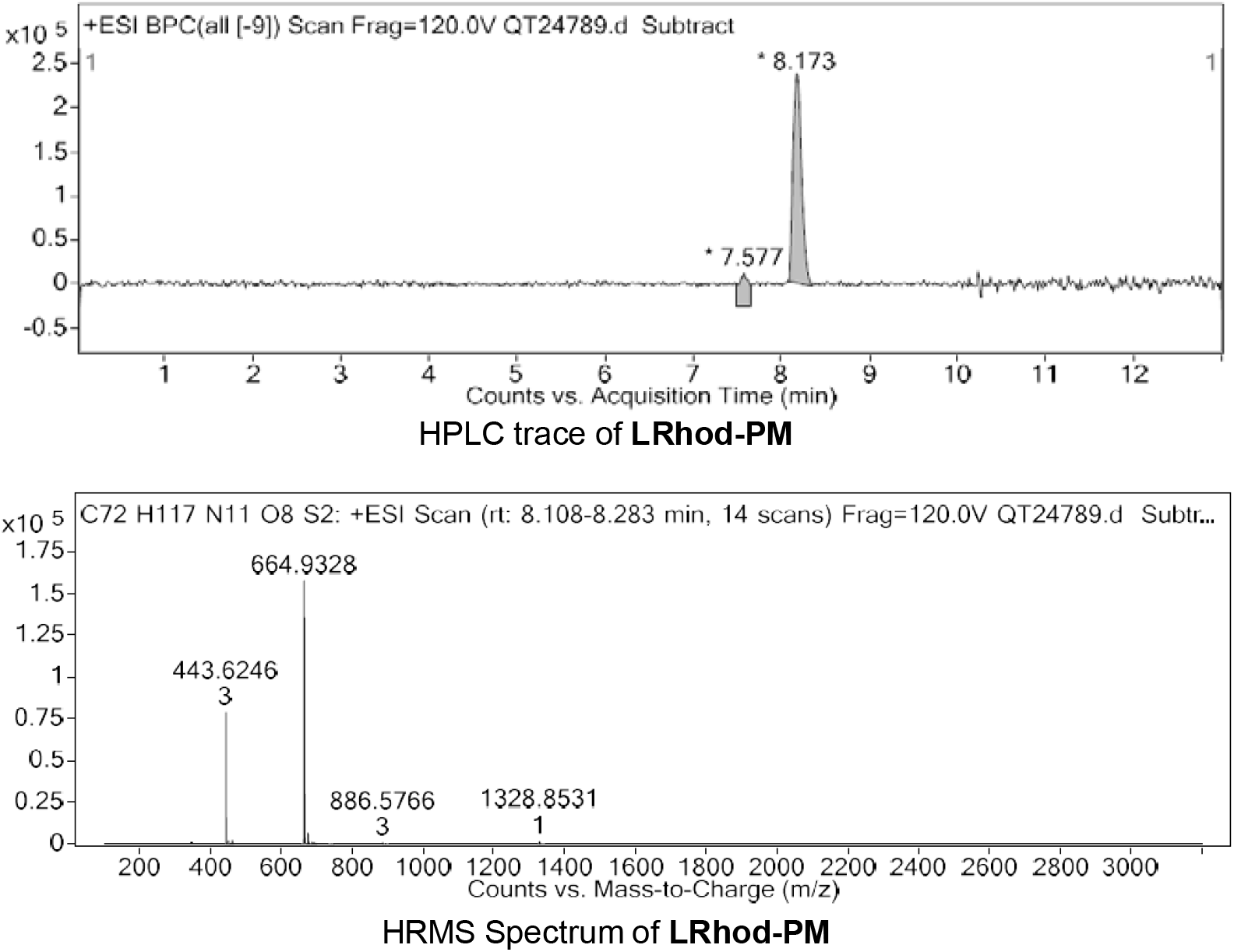

### Photooxidation studies in lipid vesicles. Method to determine kinetic constants

An analysis of the transformations occurring upon irradiation was carried out using a kinetic model based on a serial reaction mechanism. LRhod-PM is photoactivated into Rhod-PM with an associate kinetic rate constant k_Pa_, and Rhod-PM is subsequently photobleached with a kinetic rate constant k_Bl_ considering the following pathway:

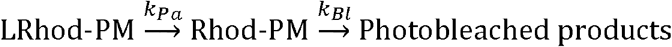

In this system, the evolution of the chemical species concentrations over time are given by the following equations:

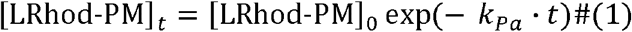

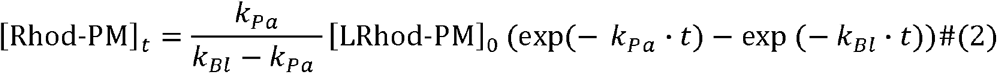

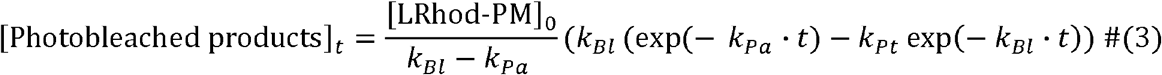

Proportionality between concentration and fluorescence intensity (FI) for low concentration samples is expressed as:^2^

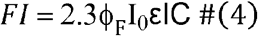

with φ_F_ : Fluorescence quantum yield, I_0_: intensity of the incident light, ε: molar-absorption coefficient of the dye at the excitation wavelength (L.mol^-1^ .cm^-1^), l: optical path length, C: concentration of the dye.

Finally, combining equation (2) and (4), kinetic rate constants k_*Pa*_ and k_*Bl*_ were determined by fitting evolution of Rhod-PM fluorescence intensity integration over time based on the following equation:

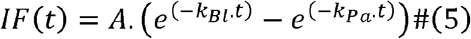

**Figure S1.**
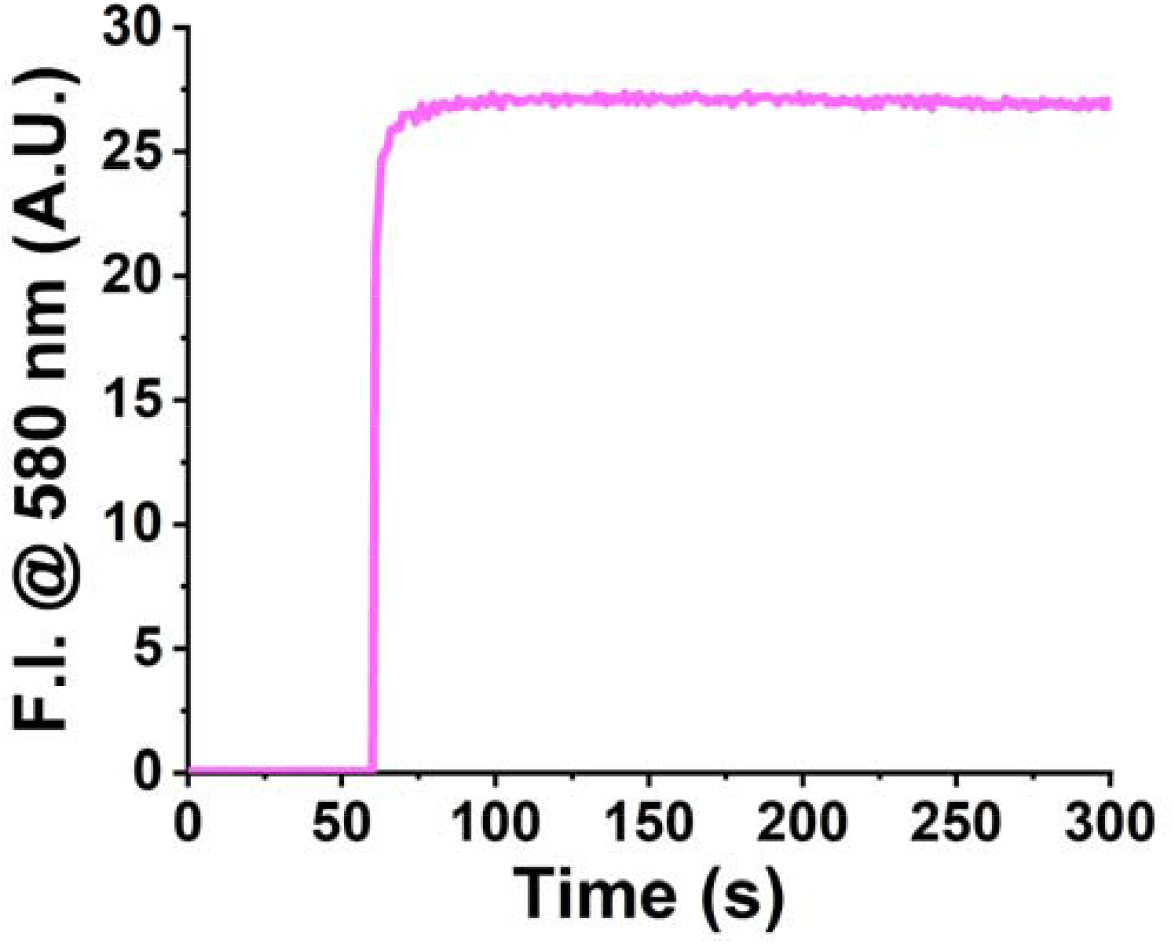
Kinetics of binding of Rhod-PM in plasma membrane model (DOPC LUVs) showing fast de-aggregation. Rhod-PM (200 nM) was added after 1 min to a solution of LUVs (lipid concentration 200 µM) and the fluorescence intensity was monitored at 580 nm.

**Figure S2.**
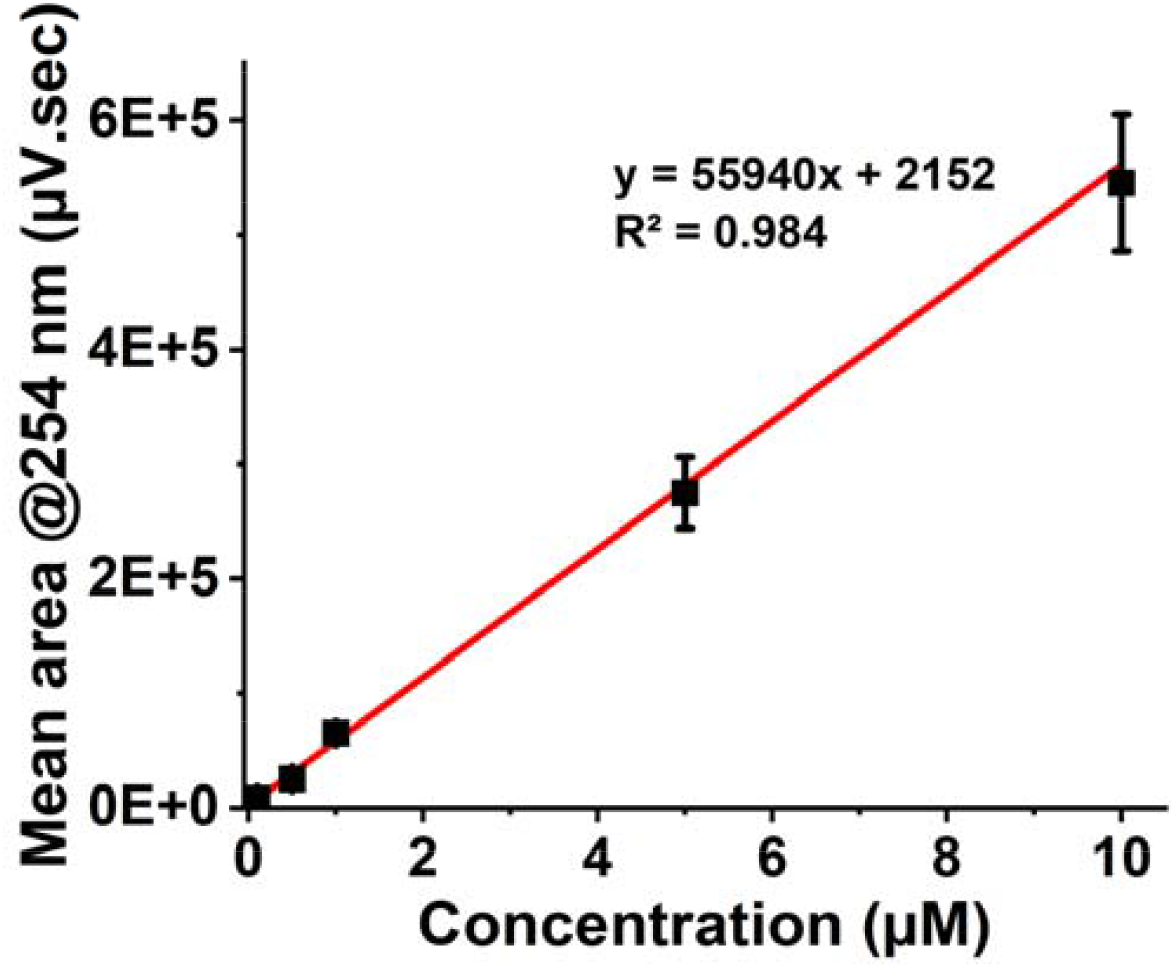
HPLC calibration curve of Rhod-DA at 254 nm. Each concentration was analyzed in triplicate.

**Figure S3.**
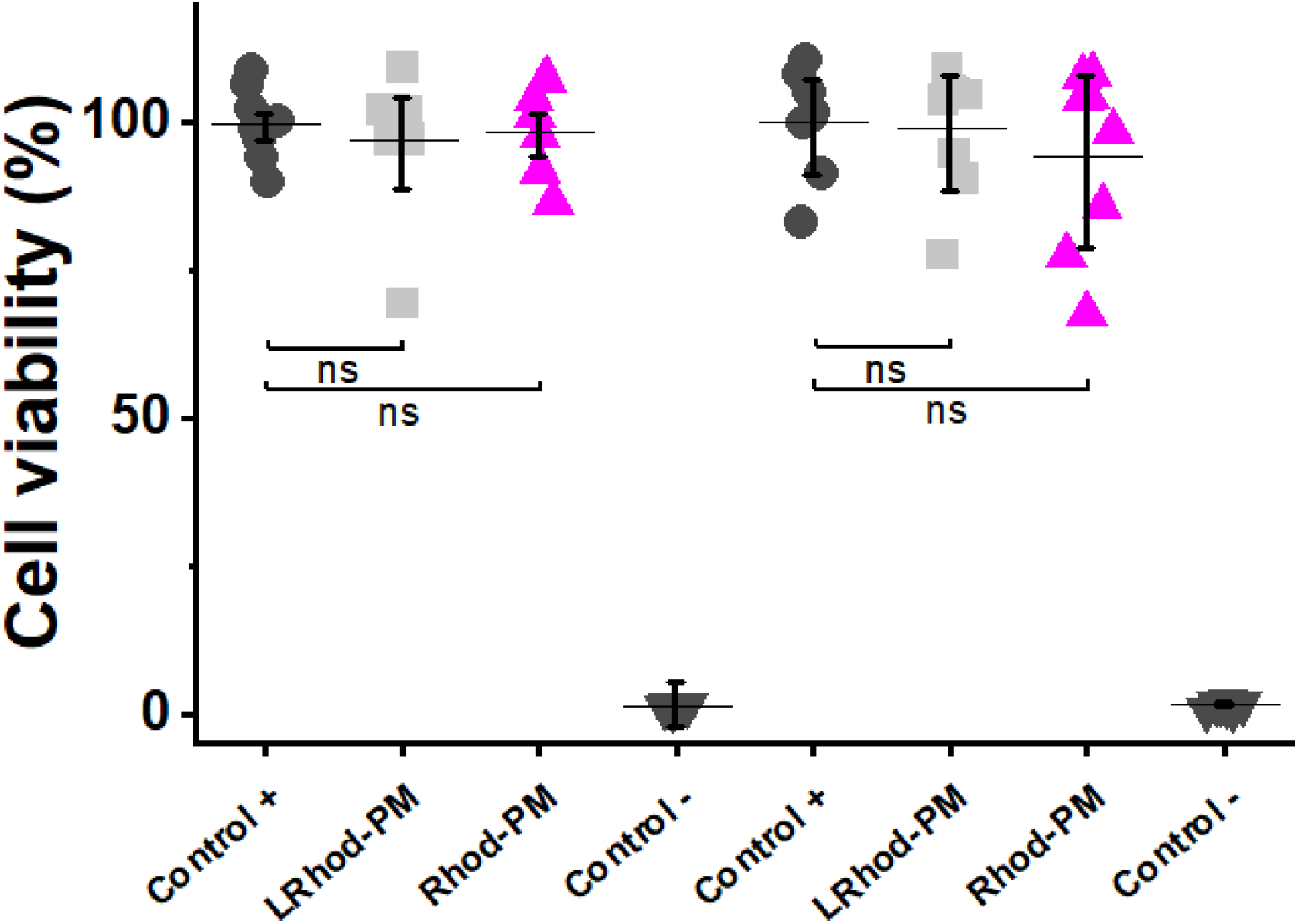
Cytotoxicity and phototoxicity assays. MTT cell viability assays in HeLa cells incubated for 30 min with Rhod-PM and LRhod-PM (200 nM) showing no sign of cytotoxicity nor phototoxicity. For cytotoxicity assay (left), the cells were not irradiated. For photocytotoxicity assay (right), the irradiation was performed by irradiating the plate 30 min using a 525 nm LED Light Source. Data are presented as mean ± standard error, n=8. ns: non-significative (a = 95%, p=0.05).

**Figure S4.**
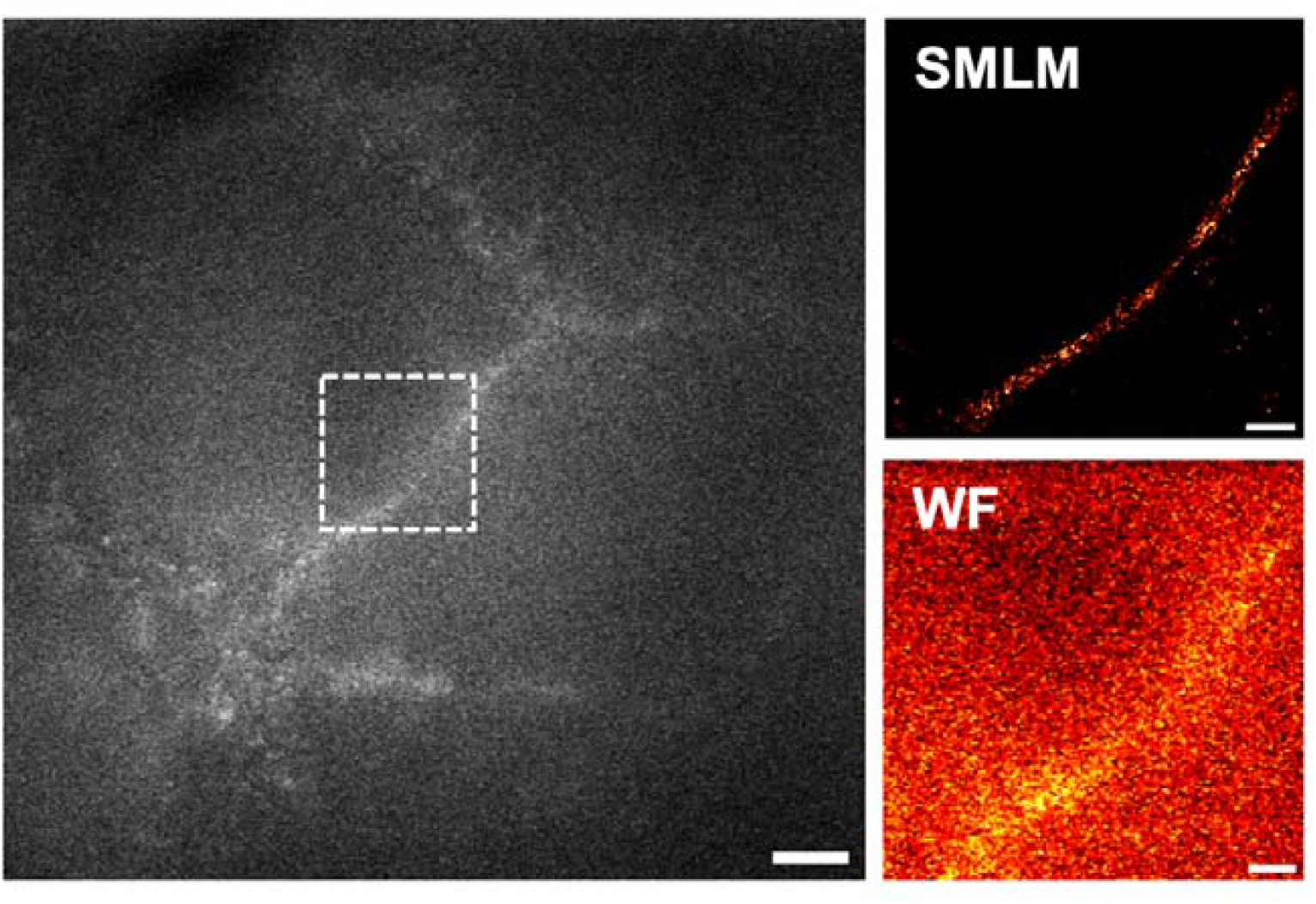
Live-cell SMLM with Rhod-PM (20 nM). Left: Widefield image after irradiation. Scale bar: 5 µm. Right (LUT Fire): Live SMLM image based on 6,000 frames and 2,500 localizations and widefield image of the region of interest (white dotted frame). Scale bar: 2 µm.

